# A FGF2-mediated incoherent feedforward loop induces Erk inhibition and promotes naïve pluripotency

**DOI:** 10.1101/2020.11.11.378869

**Authors:** Borzo Gharibi, Emanuel Gonçalves, Buhe Nashun, Alex Montoya, Katherine Mankalow, Stephanie Strohbuecker, Rahuman S M Sheriff, Alessandro Cicarrelli, Joana Carvalho, Emma Nye, Holger Kramer, Ian Rosewell, Petra Hajkova, Pedro Beltrao, Silvia D. M. Santos

**Affiliations:** The Francis Crick Institute, London, UK; Welcome Sanger Institute, Hinxton, Cambridge, UK; MRC-London Institute of Medical Sciences, Imperial College London, London, UK; School of Life Sciences, Inner Mongolia University, Hohhot, China; European Molecular Biology Laboratory - European Bioinformatics Institute, EMBL-EBI, Hinxton, CA, UK

## Abstract

Naïve pluripotency is a transient state during mammalian development that can be recapitulated indefinitely *in vitro* by inhibition of the mitogen-activated protein kinase (MAPK/Erk) signalling and activation of STAT and Wnt pathways. How Erk is inhibited *in vivo* to promote naïve pluripotency remains largely unknown. By combining live cell imaging and quantitative proteomics we found that FGF2, a known Erk activator and pro-differentiation cue, induces instead long-term Erk inhibition in both ES cells and mouse embryos. We show that Erk inhibition results from a FGF2-induced incoherent feedforward loop. Importantly, we see that FGF2 induces up-regulation of naïve pluripotency factors, down-regulation of DNA methylation by suppression of *de novo* DNA methylases thereby helping maintain naïve pluripotency. We show that FGF2 is expressed maternally and propose that integration of signals from the embryo’s niche may contribute to the generation of embryonic lineages with the right cell proportions. We suggest that feedforward regulation may play a role driving transient, reversible developmental transitions.

## Main text

During early mammalian development extrinsic signals prompt a collection of pluripotent cells in the to undergo differentiation and give rise to all cells of an adult organism. The ability of these pluripotent cells in the epiblast to self-renew and differentiate into all embryonic lineages is known as naïve pluripotency (Nichols and Smith, 2009). In vivo, naïve pluripotency is a transient state that is characterized by specific gene expression patterns (including up-regulation of Nanog, Klf2 and Klf4 transcription factors) and specific epigenetic signatures (including DNA de-methylation) and is thought to be tightly regulated by the coordinated, synergistic activities of MAPK/Erk, Wnt and Jak/STAT signalling pathways (Burdon *et al*., 1999; Nichols and Smith, 2009; Wray, Kalkan and Smith, 2010; De Los Angeles *et al*., 2015).

Naïve pluripotency can be recapitulated and indefinitely maintained *in vitro* using mouse embryonic stem (mES) cells growing in chemically defined conditions, referred to as 2iLIF, by blocking MAPK/Erk signalling and activating Wnt and Jak/STAT pathways (Ying *et al*., 2008). Cultured naïve mES cells exhibit transcriptional and epigenetic profiles resembling the cells within the epiblast *in vivo* (Leitch *et al*., 2013; Boroviak *et al*., 2015; Galonska *et al*., 2015).

The derivation of 2iLIF culture conditions has highlighted the central role of MAPK/Erk signalling in inducing exit from pluripotency and driving lineage specification. Indeed, both genetic and biochemical studies have indicated that activation of the Erk pathway is the signal that primes mES cells for differentiation (Kunath *et al*., 2007; Stavridis *et al*., 2007).

FGF signalling is thought to be at the core of triggering Erk activation and thereby lineage commitment. The resulting Erk activation is thought to drive the pluripotency-to-differentiation transition i.e. the first lineage specification in the early blastocyst and, later, in the late blastocyst stage (Frankenberg *et al*., 2011; Kang *et al*., 2013; Molotkov *et al*., 2017). Activation of Erk has been described to induce phosphorylation of Nanog, Klf2 and Klf4 pluripotency factors leading to their degradation (Kim *et al*., 2012, 2014; Yeo *et al*., 2014) and to promote rapid DNA methylation and chromatin signatures characteristic of primed state and lineage specification (Ficz *et al*., 2013; Tee *et al*., 2014). As a consequence, naïve pluripotency becomes destabilized and naïve-to-prime transition is induced in the early embryo.

mES cells growing in 2iLIF can perpetuate a state of naïve pluripotency and have been an incredibly trackable *in vitro* system to understand naïve-to-prime transition during early development. However, naïve pluripotency is only a *transient* state *in vivo*, lasting about 24h. Moreover, it has recently been shown that sustained MAPK inhibition by 2iLIF leads to sustained downregulation of methyltranferases, irreversible epigenetic changes that ultimately impair developmental potential (Choi *et al*., 2017; Yagi *et al*., 2017)

While there is overwhelming evidence that Erk signalling promotes exit from pluripotency and inhibition of Erk is important to sustain naïve pluripotency, how Erk is transiently inactivated in the epiblast *in vivo* to maintain naïve pluripotency in pre-implantation embryos remains a mystery.

## Results

### FGF2 induces a transient Erk1/2 activation followed by long-term inhibition in mES cells

We set out to investigate how Erk1/2 activity is inhibited during naïve pluripotency we tested whether developmental growth factors could rescue a state of Erk1/2 inactivation after mES cells were released from naïve pluripotency (Figure S1A). To do this, we released naïve mES cells from 2iLIF media and replaced it with basal N2B27 media in the presence or absence of canonical developmental growth factors (Figure S1A). Stimulation of naïve mES cells with N2B27 induced a sustained activation of Erk1/2, which peaked at 5 minutes and remained on for at least 60 minutes (Figure S1A).

Similarly, treatment of mES cells with N2B27 in the presence various developmental growth factors tested gave rise to a sustained, long-lasting Erk1/2 activation showing that canonical developmental factors were unable to revert mES cells back to an Erk1/2 inactive state, characteristic of naïve pluripotency.

Unexpectedly, and in sharp contrast to other FGF family members, we found that FGF2, a known Erk1/2 activator, induced a transient activation of Erk1/2 followed by a quick Erk1/2 inhibition. After an initial activation of Erk1/2 at 5 minutes, FGF2 induces a sharp decrease in Erk1/2 activity for at least 60 minutes (Figures 1A).

Importantly, we saw that FGF2 inhibited Erk1/2 in a dose-dependent manner: stimulating cells for 30 minutes with increasing concentrations showed a linear inhibition of Erk1/2 activity (Figure 1B).

**Figure 1.**
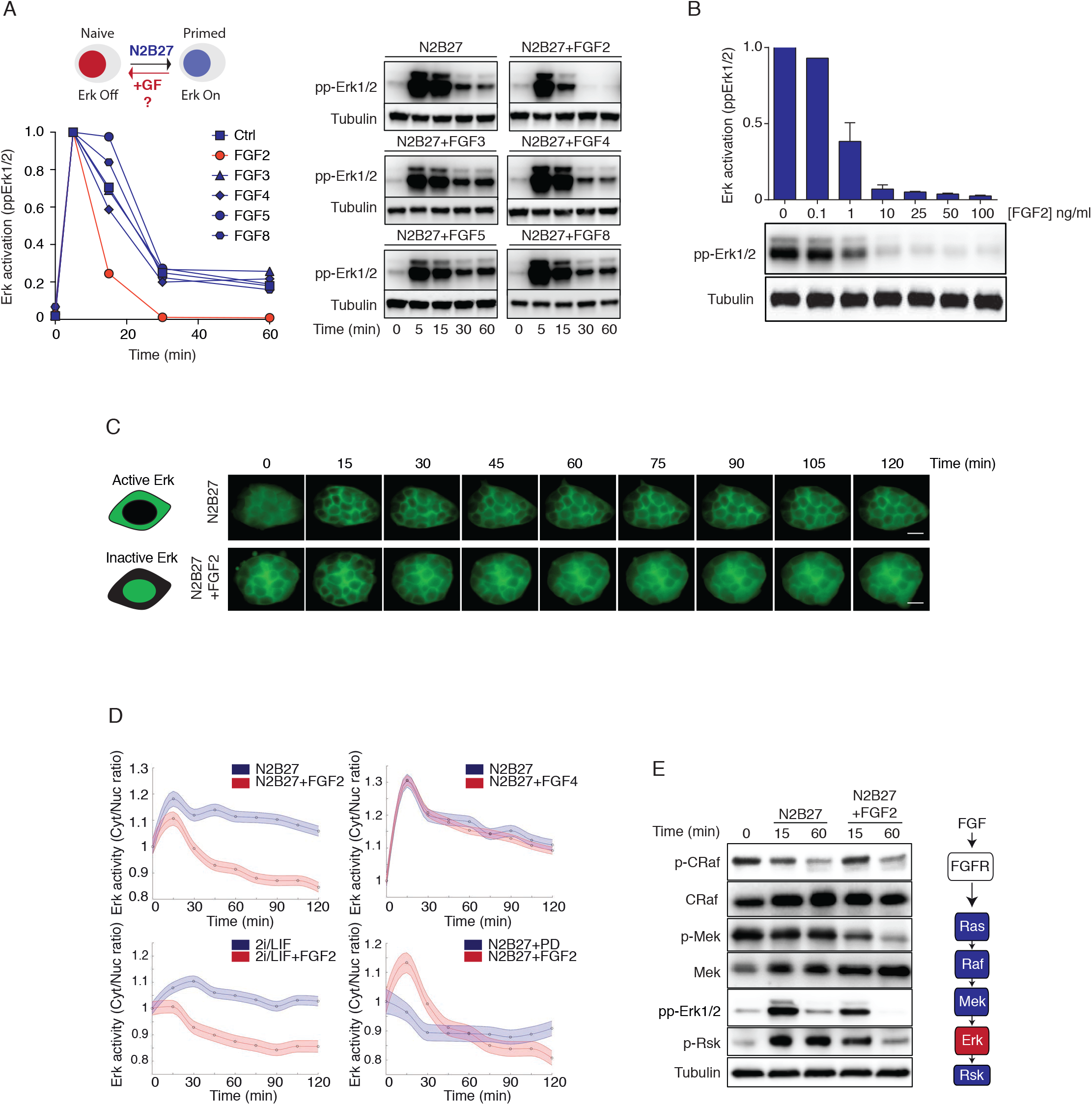
FGF2 induces transient activation and long-term inhibition of Erk1/2 in mES cells. (A) Right: Western blot time courses comparing Erk1/2 activity (pp-Erk1/2) following treatment with N2B27 and FGF family members (all at 100ng/ml). α-Tubulin was used as loading control for quantification. Representative of n=3 independent experiments. Left: Quantification of Erk1/2 activation over time for the indicated experimental conditions. (B) Dose response analysis of Erk1/2 activation (pp-Erk1/2) as a function of increasing FGF2 concentration. α-Tubulin was used as loading control for quantification. Representative of n=2 independent experiments. (C) Schematic of the Erk-KTR activity sensor used to measure Erk1 /2 activity in single live cells and representative images of a short (120mins) time course of mES cells treated with N2B27 in the presence or absence of FGF2. Scale bar represents 20μm. (D) Erk activity over 120min as measured in single cells expressing the Erk-KTR sensor treated with N2B27 in the presence or absence of FGF2 (100ng/ml), FGF4 (100ng/ml), 2i LIF or 5μM PD. C/N ratio indicates ratio of cytoplasmic to nuclear intensities. n > 100 cells were analysed for each experimental condition. (E) Western blot time course showing activation of the canonical MAPK network (C-Raf, Mek1/2, Erk1/2) and the Erk1/2 substrate Rsk1/2 following treatment with N2B27 in the presence or absence of FGF2. Schematic of the canonical FGF signalling pathway is shown on the right.

In order to confirm these observations in live cells, we established a mES stable cell line expressing an Erk1/2 activity sensor, the KTR-ERK-GFP reporter system, which has been used previously to report Erk1/2 dynamics in single cells (Regot *et al*., 2014). Erk1/2 activation was measured by the translocation of the KTR-ERK-GFP sensor from the nucleus to the cytoplasm in live cell imaging experiments (Figure 1C). Cells growing in 2iLIF were treated with N2B27 in the presence or absence of FGF2 for 2 hours. While N2B27 caused translocation of the KTR-ERK-GFP sensor to the cytoplasm (increase in Erk1/2 activity), cells treated with FGF2 showed a quick translocation to the cytoplasm followed by redistribution of the sensor to the nucleus (decrease in Erk1/2 activity) (Figure 1C). This down-regulation continued for at least 24 hours (Figures S1B, S1C and Supplemental Movie 1).

Notably, we found that FGF2 could potentiate the specific pharmacological Mek inhibitor PD032901 (PD), resulting in a synergistic long-term inhibitory effect on Erk1/2 activity when cells were treated with 2iLIF in the presence of FGF2 (Figures 1D and S1D). Using five times higher PD concentration than that in 2iLIF media confirmed after an initial short Erk1/2 activation, FGF2 induces a potent inhibition of Erk1/2 activity (Figure 1D). This is in sharp contrast to the sustained Erk1/2 activity induced in cells treated with FGF4, a growth factor which activates similar FGF receptors and has an important role in early development (Figure 1D).

FGF signalling activates the canonical Raf-Mek-Erk pathway (Lanner and Rossant, 2010). While stimulating cells with N2B27 in the presence or absence of FGF2 had no effect on total protein levels, FGF2 stimulation resulted in decreased activation levels of the whole Mek-Erk and the Erk substrate Rsk, as determined by reduced phosphorylation (Figure 1E). Interestingly, Mek activity is strikingly different in 2iLIF conditions and cells treated with FGF2.

Together these results show that FGF2 transiently activates Erk1/2 but promotes a long-term potent Erk1/2 inhibition, suggesting a unique role for FGF2 in the FGF family in regulating Erk signalling in mouse embryonic stem cells.

### FGF2 regulates Erk1/2 activity through a Ret-dependent incoherent feedforward loop

The question was thus how does FGF2 stimulation induce transient activation dynamics of Erk1/2. A simple mechanism would be that upon FGF2 stimulation Erk1/2 activates a negative regulator that would lead to down-regulation of Erk1/2 activity. Such negative feedback has been described for other growth factors (Buday L Downward J, 1995). Alternatively, FGF2 directly could down-regulate a positive regulator, or conversely, up-regulate an inhibitor of Erk1/2, a mechanism known as incoherent feedforward regulation (Alon, 2007).

In order to understand the molecular mechanism underlying FGF2 regulation of Erk1/2 dynamics we carried out a global phospho-proteomics time series analysis. Cells grown in 2iLIF media were either left untreated (control) or were treated for 15 and 60 minutes with N2B27 in presence or absence of FGF2 (Figure S2A). Three biological replicates for each time point were analysed (Figure S2B). In line with the observation that FGF2 induces Erk1/2 inhibition, volcano plots of the 9391 identified phospho-peptides show that the phospho-profiles resulting from simulating cells with FGF2 for 60 minutes were comparable to those of cells growing in 2iLIF conditions (Figure S2C).

We next performed a gene ontology (GO) analysis to identify molecular and cellular functions promoted by FGF2 stimulation (Figure 2A). The analysis highlighted that proteins involved in signal transduction differed highly between FGF2-treated and control cells (Figure 2A). As expected, a plethora of Erk1/2 substrates were differentially regulated in 2iLIF, N2B27 and N2B27+FGF2 treated cells (Figure S2D). Interestingly treating cells with 2iLIF and N2B27+FGF2 had similar effects on many Erk1/2 substrates when compared to control conditions (N2B27) (Figure S2D).

**Figure 2.**
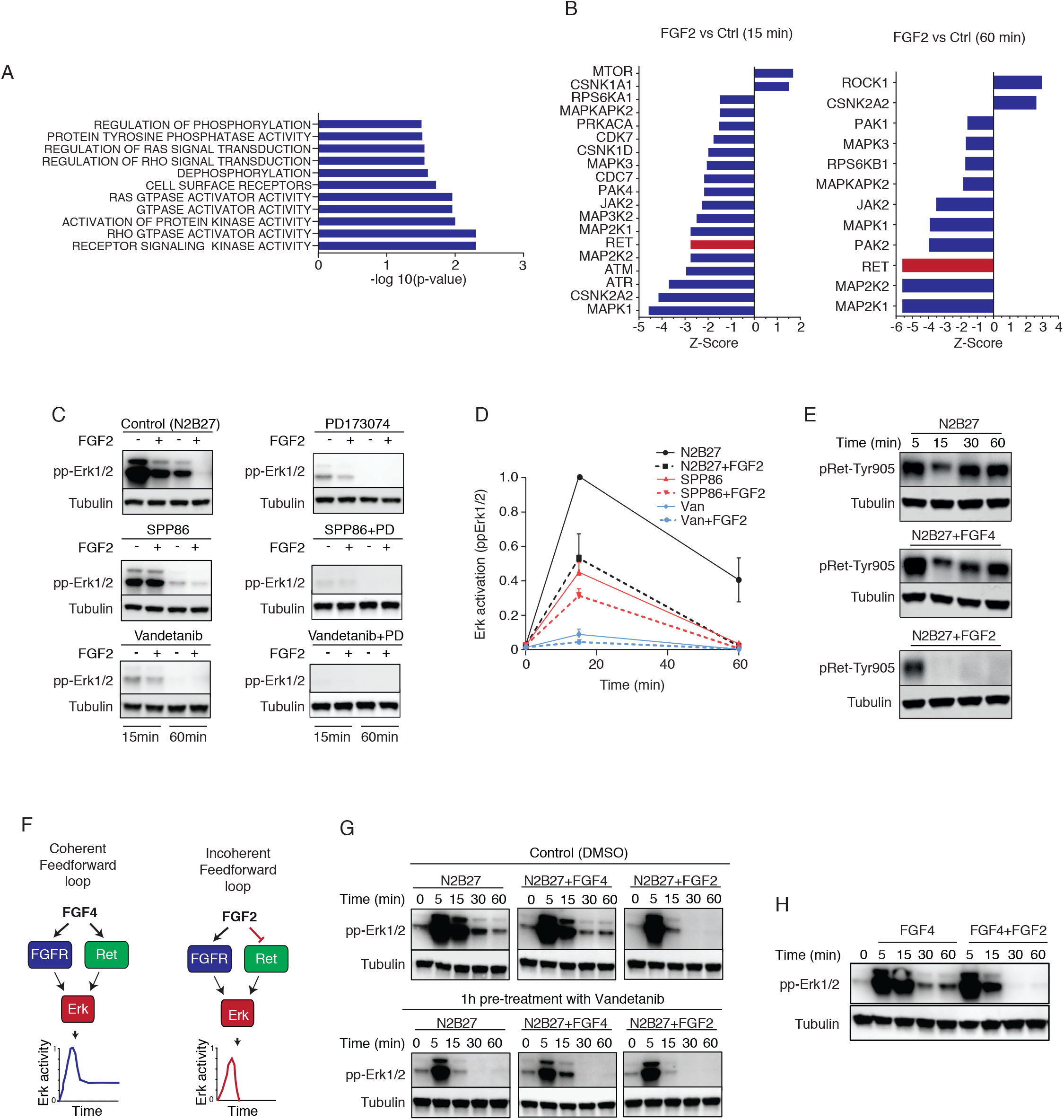
FGF2 induces a transient Erk1 /2 activation via a Ret-mediated incoherent feedforward loop. (A) Gene ontology (GO) term analysis of the differently regulated phosphopeptides for cellular functions enriched after FGF2 stimulation. (B) Kinase set enrichment analysis (KSEA) was used to predict kinases associated with FGF2 treatment at 15 and 60min. Ret is highlighted in red. (C) Western blot time courses comparing Erk1/2 activity (pp-Erk1/2) following Ret or FGFRs inhibition using SPP86 (10μM), Vandetanib (10μM) and PD173074 (1μM) in FGF2 treated or untreated cells. α-Tubulin was used as loading control for quantification. Representative of n=3 independent experiments. (D) Quantification of Erk1/2 activation over time for the experimental conditions described in D. Error bar represent mean±SD. (E) Western blot time courses comparing Ret activity (pRet-Tyr 905) in FGF2-, FGF4-treated or untreated cells. α-Tubulin was used as loading control for quantification. (F) Wiring diagram showing FGF4- and FGF2-driven coherent and incoherent feedforward loops. (G) Western blot time courses comparing Erk1/2 activity (pp-Erk1/2) following stimulation with N2B27 with either FGF2 or FGF4 in cells either left untreated or pre-treated with Ret inhibitor Vandetanib (Van) for 1hr. α-Tubulin was used as loading control for quantification. Representative of n=2 independent experiments. (H) Western blot time courses comparing Erk1/2 activity (pp-Erk1/2) following FGF4 treatment (100ng/ml) in the presence or absence of FGF2 (100ng/ml). α-Tubulin was used as loading control for quantification. Representative of n=2 independent experiments.

To further interrogate the data for possible FGF2-specific targets, we utilised a modified version of the weighted kinase set enrichment analysis (KSEA), previously used to infer kinases activities from phosphoproteomics data (Ochoa *et al*., 2016). The KSEA analysis provided a prediction of protein kinases whose activities are likely to be significantly different (up-regulated or down-regulated) when comparing N2B27+FGF2 with N2B27 treated cells (Figure 2B). We validated the top 8 predicted hits by using specific inhibitors to the different kinases and treating cells with N2B27 in the presence or absence of FGF2 (Figure S2E).

Importantly, this analysis predicted that the activity of Ret, a receptor tyrosine kinase with important roles in development (Pachnis, Mankoo and Costantini, 1993; van Weering and Bos, 1998), was significantly down-regulated in FGF2-treated cells (Figure 2B). And, notably, we found that treating cells with Ret inhibitors resulted in a striking down-regulation of Erk1/2 and mimicked the effects of FGF2 (Figures 2C, 2D). Indeed, stimulating N2B27-treated cells with three different Ret inhibitors (Vandetanib, SPP86 and Cabozantinib) showed down-regulation of Erk1/2 activity comparable to that of cells treated with FGF2. This implies that Ret is likely to act directly downstream of FGF2 (Figures 2C, 2D and S2F).

Furthermore, blocking both FGF receptor (FGFR) and Ret resulted in a synergistic effect whereby treating cells with both inhibitors completely abolished Erk1/2 activation (Figure 2C and 2D). And importantly, we see that FGF2, induces rapid inhibition Ret activity (Figure 2E). This shows that both FGF and Ret signalling are necessary for full Erk1/2 activation (Figure 2F).

Together these results suggest that FGF2 may induce Erk1/2 transient activation and long-term inhibition by a Ret-mediated incoherent feed-forward loop (Figure 2F).

To further confirm this feedforward regulation, cells were pre-treated with Ret inhibitor Vandetanib for one hour and were then stimulated with N2B27 with either FGF2 or FGF4 (Figure 2G). We observed that inhibiting Ret in FGF4-treated cells induces a transient Erk1/2 activation, essentially rewiring the Erk1/2 response to mirror that of FGF2 stimulation (Figures 2G). Similar results were obtained using a different Ret inhibitor (Figure S2G).

We explored the mechanism by which FGF2 might down-regulate Ret signalling and found that inhibiting endocytosis sustained Erk activation in FGF2-treated cells (Figure S2H). This suggests that FGF2 may promote endocytosis of the Ret receptor. In agreement with this, our phospho-proteomics analysis also revealed that FGF2 treatment elicits changes in phosphorylation of key endocytosis regulators, including Dynamin, Rab11, Rab3 and Cortactin (Figure S2I). Importantly, we see that FGF2 leads to internalization of activated Ret (Figure S2J). This confirms that FGF2 leads to internalization and down-regulation of Ret.

If the mechanism of the incoherent feedforward regulation is by FGF2-mediated endocytosis and inhibition of Ret, the prediction would be that FGF2 could override FGF4-induced sustained activation of Erk1/2. To test this, we treated cells with N2B27 with either FGF4 alone or FGF4 and FGF2 combined. We indeed see that while Erk activity was high in ES cells treated with FGF4, when FGF2 and FGF4 were combined, Erk activity became transient, resembling the response to FGF2 treatment (Figure 2H).

Taken together these data suggest a unique role for FGF2 in long-term Erk inhibition, involving a Ret-mediated incoherent feedforward regulation.

### FGF2 maintains naïve pluripotency in mES cells

We next investigated the functional consequence of Erk1/2 long-term inhibition by FGF2. The GO-term analysis performed to identify cellular functions promoted by FGF2 stimulation also highlighted significant changes in expression of pluripotency and epigenetic regulators in FGF2-treated cells (Figure S3A). Analogous to what we observed for Erk1/2 targets, 2iLIF and FGF2 culture conditions displayed very similar phosphorylation patterns of pluripotency regulators (Figure S3B). This raised the possibility that FGF2 might have an important role in regulating pluripotency.

To test this, we first checked how the expression of the core pluripotency factors (Nanog, Sox2 and Oct4) changed in the presence of FGF2. Cells growing in 2iLIF were treated with N2B27 in the presence or absence of FGF2 for 48 hours (Figure 3A). As expected, control cells treated with N2B27 showed a decrease in Nanog and Sox2 protein levels. Remarkably, FGF2 prevented this and, instead, maintained significantly higher levels of both pluripotent genes (Figures 3A, 3B). There was no observable effect of FGF2 on Oct4 (Figures 3A, 3B). This effect of FGF2 on pluripotent factors was also replicated in the E14 mES cells, showing that this effect is not specific to a particular mES cell line (Figures S3C, S3D).

**Figure 3.**
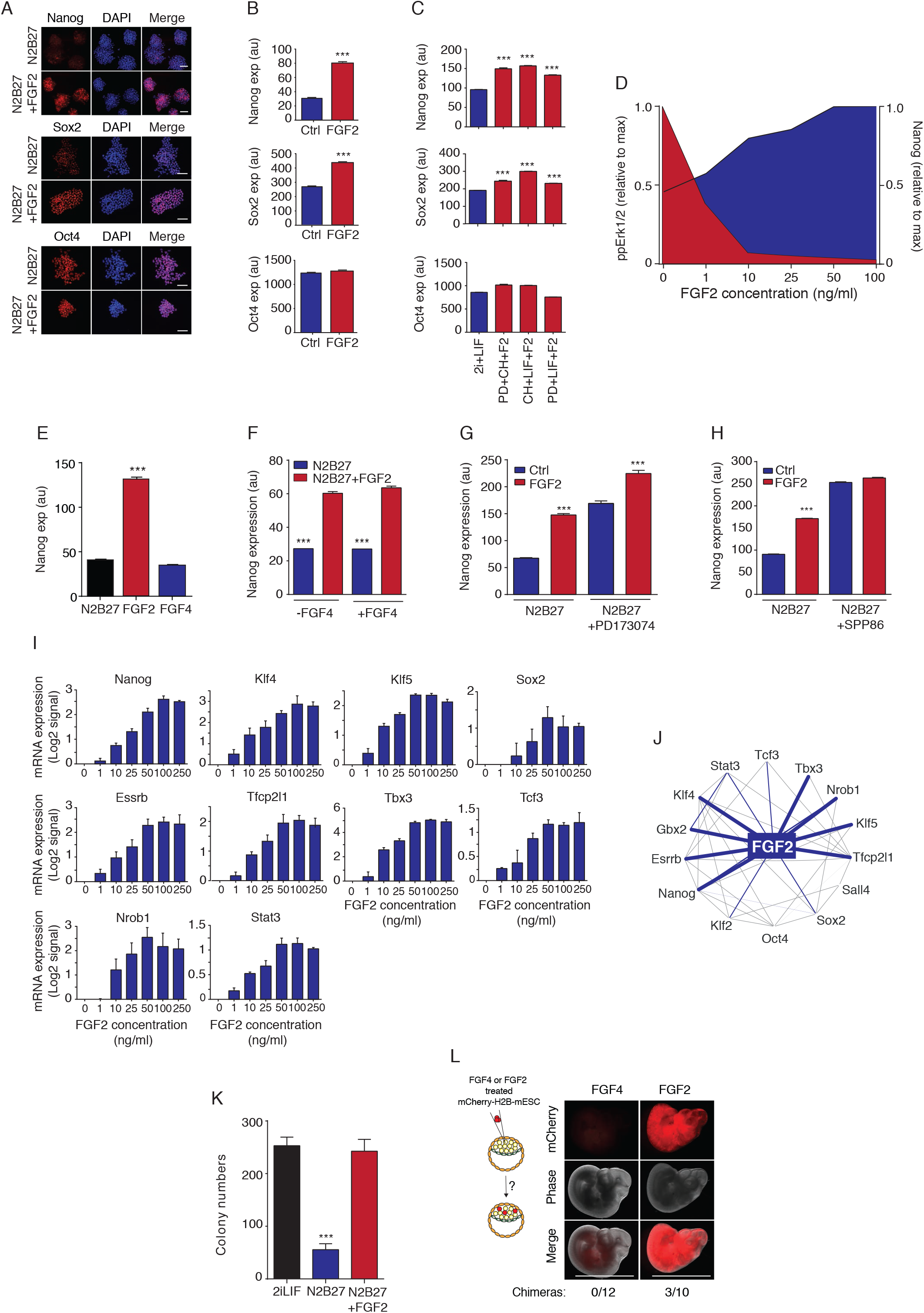
FGF2 supports naïve pluripotency in mouse ES cells. (A) Representative immunofluorescent (IF) images of the core pluripotent factors Nanog, Sox2 and Oct4 in cells treated with N2B27 in the presence or absence of FGF2 (100ng/ml) for 48h. Scale bar represents 50μm. (B) Quantification of Nanog, Sox2 and Oct4 protein expression at 48h in the experimental conditions described in A. n>500 cells were analyzed for each experimental condition. The error bars show mean ±SEM. *** p<0.0001 by KS- and Mann Whitney tests. n>3 independent experiments. (C) Quantification of protein levels of core pluripotency factors Nanog, Sox2 and Oct4 by IF after five passages in cells treated with different combination of PD, CH, LIF and FGF2. n>500 cells were analyzed for each experimental condition. The error bars show mean ±SEM. *** p<0.0001. (D) Dose response curves of Erk activity (pp-Erk1/2) and Nanog protein expression to increasing FGF2 concentrations. (E) Nanog expression following treatment with FGF2 (100ng/ml) or FGF4 (100ng/ml) for 24 hours. n > 500 cells were analyzed for each experimental condition. The error bars show mean ±SEM. *** p<0.0001. (F) Nanog expression in cells cultured for 48h in N2B27 and FGF4 in presence or absence of FGF2 (100ng/ml). n>500 cells were analysed for each experimental condition. The error bars show mean ±SEM. *** p<0.0001. (G) Nanog expression following treatment with FGF2 (100ng/ml) in presence or absence of FGFRs inhibitor PD173074 (1μM). n > 500 cells were analysed for each experimental condition. The error bars show mean ±SEM. *** p<0.0001. (H) Nanog expression following treatment with FGF2 (100ng/ml) in presence or absence of Ret inhibitor SPP86 (10μM). n > 500 cells were analysed for each experimental condition. The error bars show mean ±SEM. *** p<0.0001. (I) Expression of naïve pluripotency associated transcription factors in cells cultured for 48hrs in N2B27 with increasing concentrations of FGF2. Data was normalized to house-keeping gene GAPDH and shown as log2 fold change relative to control N2B27-treated cells, which was set as 0. The error bars show mean ±SD for n=3 independent experiments. (J) Transcriptional regulatory network regulated by FGF2 (based on the data shown in S3I) and the possible interaction between pluripotency transcription factors based on previously published networks^32^. Interaction is represented by a blue line; the thickness of each line reflects the degree of stimulation by FGF2. (K) Clonogenicity of ES cells cultured in 2i LIF or N2B27 in the presence or absence of FGF2 for 48 hours. The error bars show mean ±SD, *** p<0.0001 for n=3 independent experiments. (L) Representative images of E9.5 chimeras generated by injection of B6N6-H2B-mCherry mES cells treated with N2B27+FG2 (100ng/ml) or N2B27+FGF4 (100ng/ml) into E4.5 C57BL/6 blastocysts. mCherry images show chimeric contribution. 3 out of 10 and 0 out of 12 dissected embryos injected with FGF2 and FGF4 treated cells, respectively, resulted in chimeras. Scale bar represents 2.5mm.

We next generated a stable Nanog-2A-mCherry CRISPR knock-in mES line to study naïve pluripotency in live single cells. In mES cells, naïve pluripotency is marked by a uniform and elevated expression of Nanog (Nichols and Smith, 2009; Nichols *et al*., 2009; Silva *et al*., 2009; Muñoz Descalzo *et al*., 2012).

Cells cultured in 2iLIF media for 48 hours formed tight colonies and showed uniform Nanog expression. Withdrawal of any component of 2iLIF resulted in a significant reduction in Nanog expression (Figures S3E and S3F). Surprisingly, adding FGF2 to any of these culture conditions reverted Nanog down-regulation and led to an increase in Nanog expression in all conditions (Figures S3E and S3F). Consistently, addition of FGF2 gave rise to morphology characteristic of naïve pluripotency, with tighter, more spherical colonies with a compacted centre (Figures S3F). As seen before, while Sox2 protein levels were also affected by FGF2 treatment, Oct4 expression remained relatively unchanged (Figures S3E).

To explore whether FGF2 is able to support long-term maintenance of naïve pluripotency, we replaced either CH, PD or LIF with FGF2 for five passages (Figure 3C). We observed that FGF2 could indeed substitute for any of the components in 2iLIF media and maintain high levels of Nanog and Sox2 expression for five passages, thereby maintaining naïve pluripotency long-term (Figures 3C).

Interestingly, we see that the effect of FGF2 in Nanog is dose dependent and we observed an inverse correlation between Erk1/2 activity and Nanog expression to FGF2 concentration (Figure 3D). Cells stimulated with increasing doses of FGF2 showed downregulation of Erk1/2 activity and up-regulation of Nanog expression (Figures 3D). A similar up-regulation of Nanog protein was also seen when treating cells with increasing concentrations of the Erk1/2 inhibitor PD (Figure S3G), confirming that FGF2 behaves as an inhibitor of Erk1/2.

In sharp contrast with these observations, stimulating cells with N2B27 in the presence of FGF4 showed down-regulation of Nanog (Figures 3E). This could be rescued by combining both FGF4 and FGF2 (Figure 3F), confirming that FGF4 and FGF2 have opposing roles on Nanog expression and, thereby, on pluripotency.

In line with what was observed for Erk1/2 dynamics, maintenance of Nanog expression by FGF2 depends on both activation of FGFR and down-regulation of Ret signalling as inhibiting either FGFR or Ret with specific inhibitors promotes Nanog up-regulation (Figures 3G and 3H).

Finally, we investigated further how FGF2 impacts on the expression of several known ancillary pluripotency factors. Cells grown in 2iLIF and subjected to N2B27 stimulation for 48 hours with increasing doses of FGF2 showed up-regulation of all canonical pluripotency factors tested (Figures 3I). These included canonical naïve pluripotency markers such as Klf4, Klf5, Tfcp2l1, Nrob1 and Essrb. Merging our gene expression data with published data on transcriptional interactions in pluripotency gave rise to a FGF2-pluripotency factors regulatory network (Figure 3J). These interactions were confirmed by monitoring the expression of pluripotency transcription factors in culture conditions consisting of FGF2 and either CH, PD or LIF (Figure S3H). We indeed found that 4 factors (Nanog, Tfcp2l1, Tbx3 and Gbx2) were consistently expressed (Nanog, Tfcp2l1, Tbx3) or repressed (Gbx2) in all conditions containing FGF2, suggesting that they might mediate the effect of FGF2 in naïve pluripotency (Figure S3H).

We further tested the role of FGF2 in naïve promoting pluripotency by performing a clonogenicity assay and we saw that the capacity for self-renewal (as measured by colony formation) was very similar in 2iLIF conditions and in FGF2-treated cells (Figure 3K). It is worth noting that measuring the time between two consecutive mitoses revealed no measurable differences in cell cycle length between cells treated with N2B27 and N2B27+FGF2 (Figure S3I).

Finally, we tested whether FGF2 treated mES cells could re-enter development and contribute to the generation of mouse chimeras when introduced into host embryos, as expected if FGF2 supports naïve pluripotency (Figure 3L). B6N6-H2B-mCherry mES cells treated with either FGF2 or FGF4 were injected into E4.5 blastocysts and transferred to pseudo-pregnant recipient mice. Embryos were dissected at E9.5 and assessed for chimeric contribution. Strikingly, we see that while no chimeric contributions were seen for FGF4, FGF2 treated cells contributed to generation of chimeras (Figure 3L). This data strongly supports that FGF2 maintains naïve pluripotency in mES cells.

Taken together, these results show that FGF2 up-regulates expression of Nanog and other canonical pluripotency regulators and is likely to have a direct role in both inducing and maintaining naïve pluripotency in mES cells.

### FGF2 suppresses *de novo* methyltransferases and DNA methylation

One of the hallmarks of naïve pluripotency is genome-wide hypomethylation and transcriptional changes in methyltransferases (Leitch *et al*., 2013). We tested whether FGF2 induced any changes in the expression levels of methyltransferases and of DNA methylation patterns.

While levels of Dnmt1 and of two members of the Ten eleven translocation (Tet) family of enzymes, Tet1 and Tet3 remained relatively unchanged (Figure 4A), saw that cells cultured in N2B27 in the presence of FGF2 had significantly lower mRNA and protein levels of the *de novo* methyltransferases Dnmt3a and Dnmt3b (Figures 4A and 4B). These were long-term changes, as Dnmt3a and Dnmt3b remained down-regulated after culturing cells for five passages in 2iLIF conditions, substituting the Erk1/2 inhibitor PD for FGF2 (Figure S4A). In fact, FGF2 treatment outperforms Erk1/2 inhibition by PD down-regulating these methyltransferases (Figure S4A).

**Figure 4.**
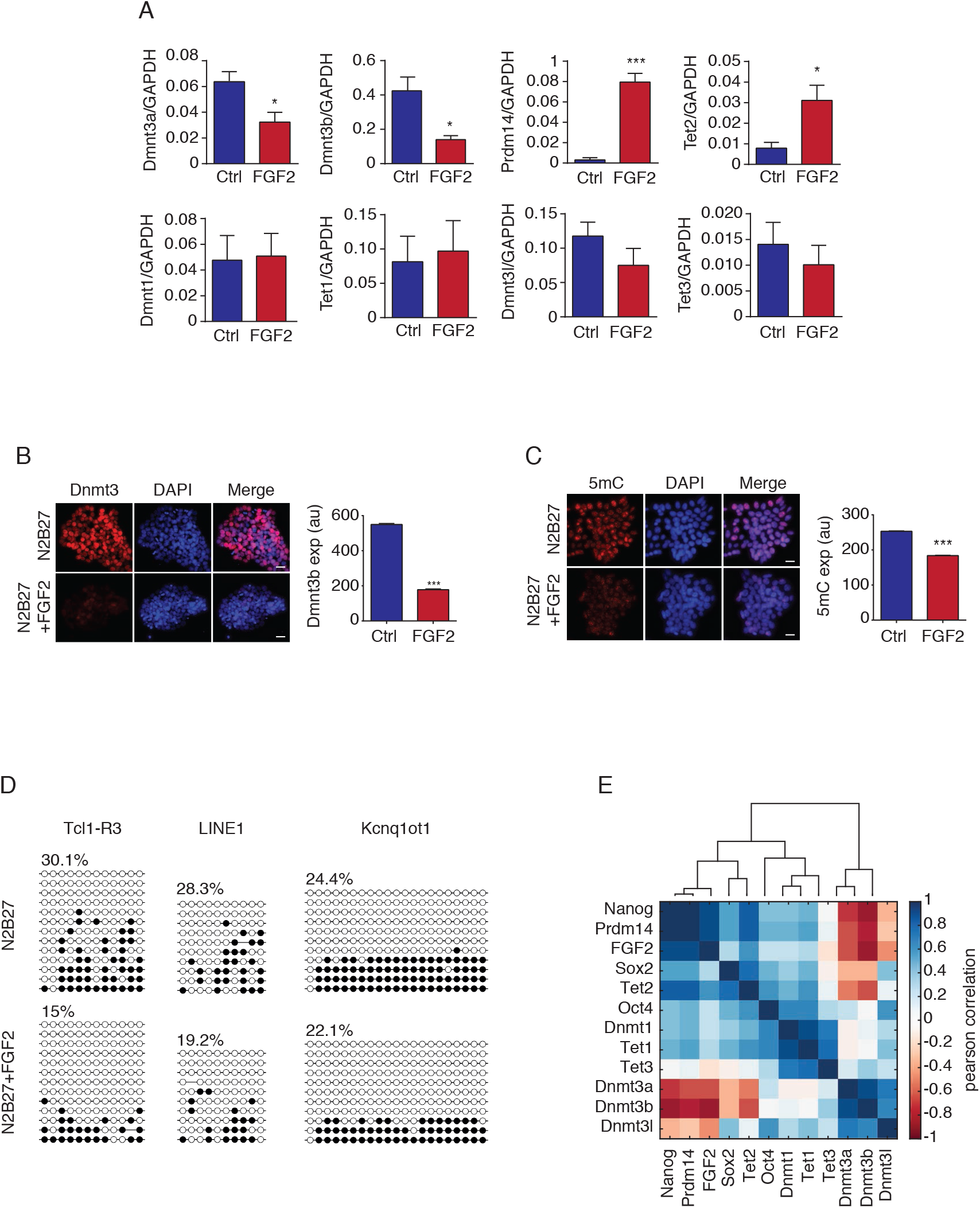
FGF2 supports naïve pluripotency in mouse ES cells by suppressing *de novo* DNA methylation. (A) Gene expression levels of epigenetic regulators in cells cultured in N2B27 in the presence or absence of FGF2 (100ng/ml). Data was normalized to the house-keeping gene GAPDH. n = 3 independent experiments. The error bars show mean ±SD. *** p<0.0001 (B) Expression and quantification of Dnmt3 protein levels in cells cultured in N2B27 in the presence or absence of FGF2. n>500 cells were analysed for each experimental condition. The error bars show mean ±SEM. *** p<0.0001. Representative images are shown in the left. Scale bar represents 20μm. (C) Expression and quantification of 5mC levels in cells cultured in N2B27 in the presence or absence of FGF2. n>500 cells were analysed for each experimental condition. The error bars show mean ±SEM. *** p<0.0001. Representative images are shown on the left. Scale bar represents 20μm. (D) Bisulfite sequencing analysis of DNA methylation levels for Tcl1-R3, LINE1and Kcnq1ot1 in cells cultured for 48h in N2B27 in the presence or absence of FGF2. White and black circles represent the unmethylated and methylated CpG sites, respectively. Level of methylation is shown as percentage. (E) Correlation analysis between expression levels of pluripotency factors, and epigenetic regulators after FGF2 stimulation. Pearson correlation coefficients were reverse engineered from gene expression changes in Nanog and Sox2 perturbed cells treated with FGF2. Scale represents Person correlation coefficients.

Recently it has been reported that that the transcriptional regulator Prdm14 is involved in the suppression of Dnmt3a and Dnmt3b and activation of Tet hydroxylases during the 2i-induced hypomethylation in naïve ES cells (Leitch *et al*., 2013; Yamaji *et al*., 2013; Okashita *et al*., 2014). In line with this, we found that cells grown in presence of FGF2 showed a higher level of Prdm14 and of the Tet2 DNA hydroxylase (Figure 4A and Figure S4A).

We next assessed whether these changes in expression of de *novo* methyltransferases lead to changes in DNA methylation. We found that FGF2 induced a reduction in global 5mC content suggesting that FGF2 induces to global hypomethylation (Figure 4C). By performing bisulfide sequencing, we quantified the level of DNA methylation on the repetitive sequences long interspersed nuclear element 1 (LINE1) promoter and the previously defined region 3 around the transcription start sites of Tcl-1 promoter. Both Tcl-R3 and LINE1 are hypomethylated in naïve pluripotent state cells following a Prdm14-driven epigenetic re-configuration (Yamaji *et al*., 2013). Strikingly, we found that FGF2-treated cells showed a considerably lower level of methylation for Tcl1-R3 (30.1% versus 15%) and LINE1 (28.3% versus 19.2%) (Figure 4D). In contrast, bisulfite sequencing revealed that methylation is relatively unchanged at imprinted differentially methylated region Kcnq1ot1 (22.1% versus 24.4), which is known to be regulated by Dnmt1 (Figure 4D).

Finally, we used a reverse engineering approach to infer FGF2-induced relationships between pluripotency and methylation (Figures 4E and S4B). We knocked down Nanog and Sox2 by shRNA and compared expression levels of pluripotency factors and epigenetic regulators in control and perturbed cells treated with FGF2. Pearson coefficient was used as a metric to quantify relationships amongst different genes (Figures 4E and S4B). This approach confirmed that FGF2 and naïve pluripotency factors Nanog and Prdm14 all clustered together and their expression is inversely correlated with that of *de novo* methyltransferases and highlighted the impact of FGF2 in promoting naïve pluripotency.

Taken together our data suggest that FGF2 leads to loss of *de novo* methyltransferases and DNA methylation, thereby promoting naïve pluripotency.

### FGF2 effect on pluripotency is time-dependent

So far, we have shown that FGF2 maintains pluripotency when cells growing in 2iLIF are induced to leave naïve pluripotency by N2B27 treatment. However, a wealth of research has also shown that FGF2 maintains pluripotency in the post-implantation epiblast (EpiSC) (Brons *et al*., 2007). FGF2 is actually not expressed in EpiSC and exogenous FGF2 seems to have little effect on expression of Nanog in these cells(Greber *et al*., 2010). We therefore sought to explore whether there is a developmental time window during which FGF2 can maintain naïve pluripotency.

In order to do this, naïve mES cells were primed by removal of 2iLIF and grown in N2B27 for 48 hours. This led to a gradual reduction in expression of Nanog in control cells treated with N2B27 alone (Figure 5A and S5A). To test whether higher levels of Nanog expression could be restored after priming and naïve pluripotency rescued, FGF2 was added at 3, 6, 12, 24 or 48 hours after N2B27 stimulation for further 24 hours (Figure 5A and S5A). We found that there is a window of time (24 hours after priming) where FGF2 is able to revert cells the loss of Nanog expression and bring cells back to a more pluripotent state (Figures 5A and S5A). This window of reversibility is lost after 24 hours of N2B27 exposure where treatment with FGF2 can no longer sustain high Nanog levels (Figures 5A and S5A).

**Figure 5.**
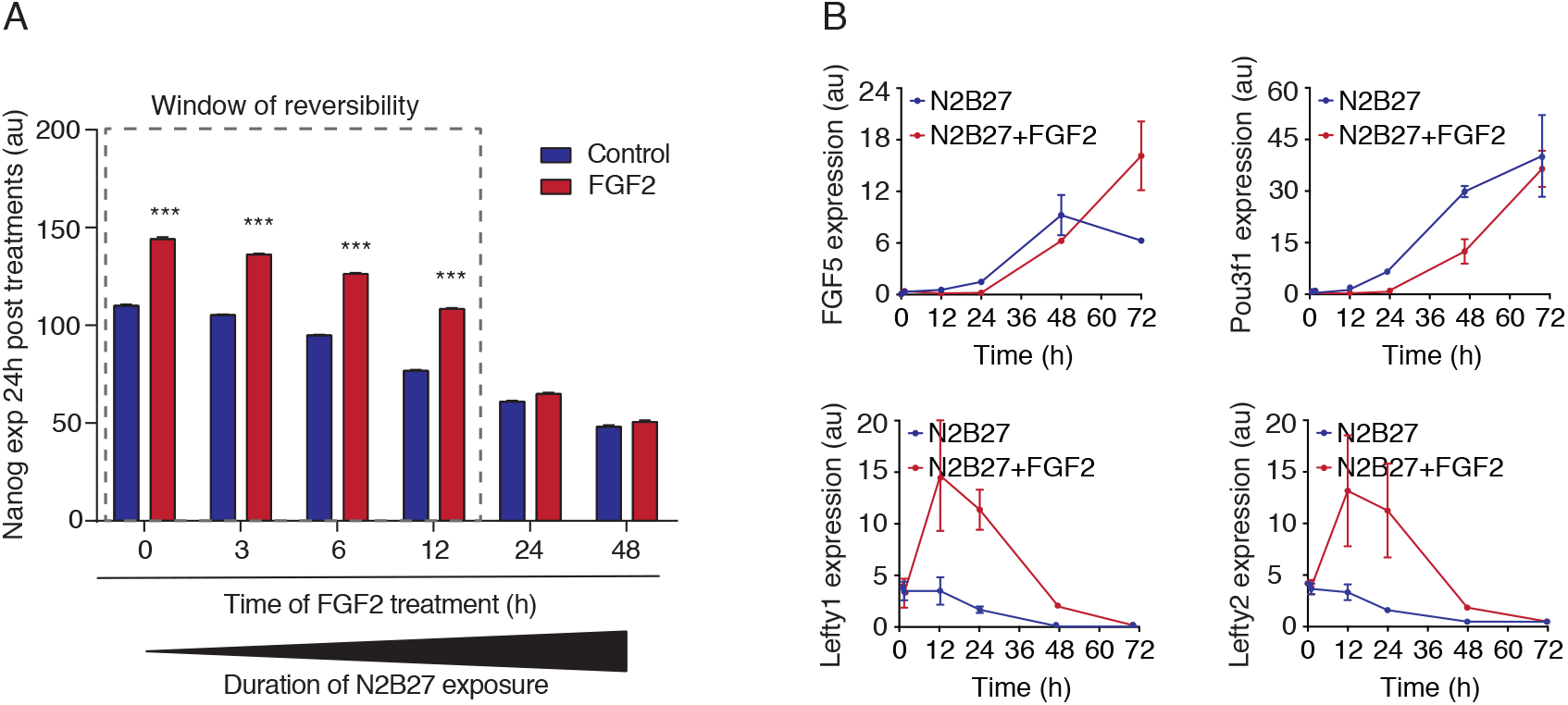
FGF2 rescues high Nanog expression for up to 24h following N2B27 treatment. (A) Nanog expression after switching cells from naïve pluripotency (2iLIF) to N2B27 followed by stimulation with FGF2 (100ng/ml) at 3, 6, 12, 24 and 48 hours or left untreated. Nanog protein levels were measured 24h after FGF2 addition at each time point. Window of FGF2 reversibility to pluripotent state is highlighted. n>500 cells were analyzed for each experimental condition. The error bars show mean ±SEM. (B) Gene expression levels of primed pluripotency regulators in cells cultured in N2B27 in the presence or absence of FGF2 (100ng/ml). Data was normalized to the house-keeping gene GAPDH. n=3 independent experiments. The error bars show mean ±SD.

In line with the idea that FGF2 promotes naïve pluripotency in a time-dependent manner, we see that canonical primed pluripotency specific genes such as FGF5 and Pou3f1 are upregulated only after 24 hours of FGF2 stimulation (Figure 5B), at which time, the activin/nodal inhibitors Lefty1 and Lefty2 also decrease their expression (Figure 5B).

These observations suggest that the potential role of FGF2 in maintaining pluripotency is limited to the naïve state (potentially corresponding to late blastocyst stage *in-vivo)* but that it does not support pluripotency in primed cells (corresponding to the pre-gastrulation embryo).

### FGF2 inhibits Erk1/2 activity and helps maintain naïve pluripotency in late blastocysts

We next questioned whether these observations were true *in vivo*, in the early embryo.

We started by examining the effect of FGF2 on the Erk activity. We isolated mouse embryos at E2.5 and developed them into E4.5 embryonic stage by *ex-vivo* culture in 2iLIF media. This has been shown to generate functional epiblast cells presenting ground state pluripotency characteristics (Nichols and Smith, 2009). At this stage, embryos cultured in 2iLIF displayed a complete ablation of Erk1/2 activity, as seen by cytoplasmic staining of a specific phospho-Erk1/2 antibody (ppErk) (Figure 6A). On the contrary, embryos treated with N2B27 in the presence or absence of FGF4 for 30 minutes caused a significant increase in Erk1 /2 activity, as seen by nuclear translocation of phosphorylated Erk1/2 (Figure 6A). ppErk nuclear accumulation was blocked by treatment of FGF4 in the presence of a Mek1/2 inhibitor (PD), showing that the staining is specific for Erk1/2 activity. Notably, in line with our findings in ES cells, treating embryos FGF2 for 30 minutes blocked Erk1/2 activation (Figure 6A). Similar results were seen using an antigen-retrieval method for immune-fluorescence (Figure S6A).

**Figure 6.**
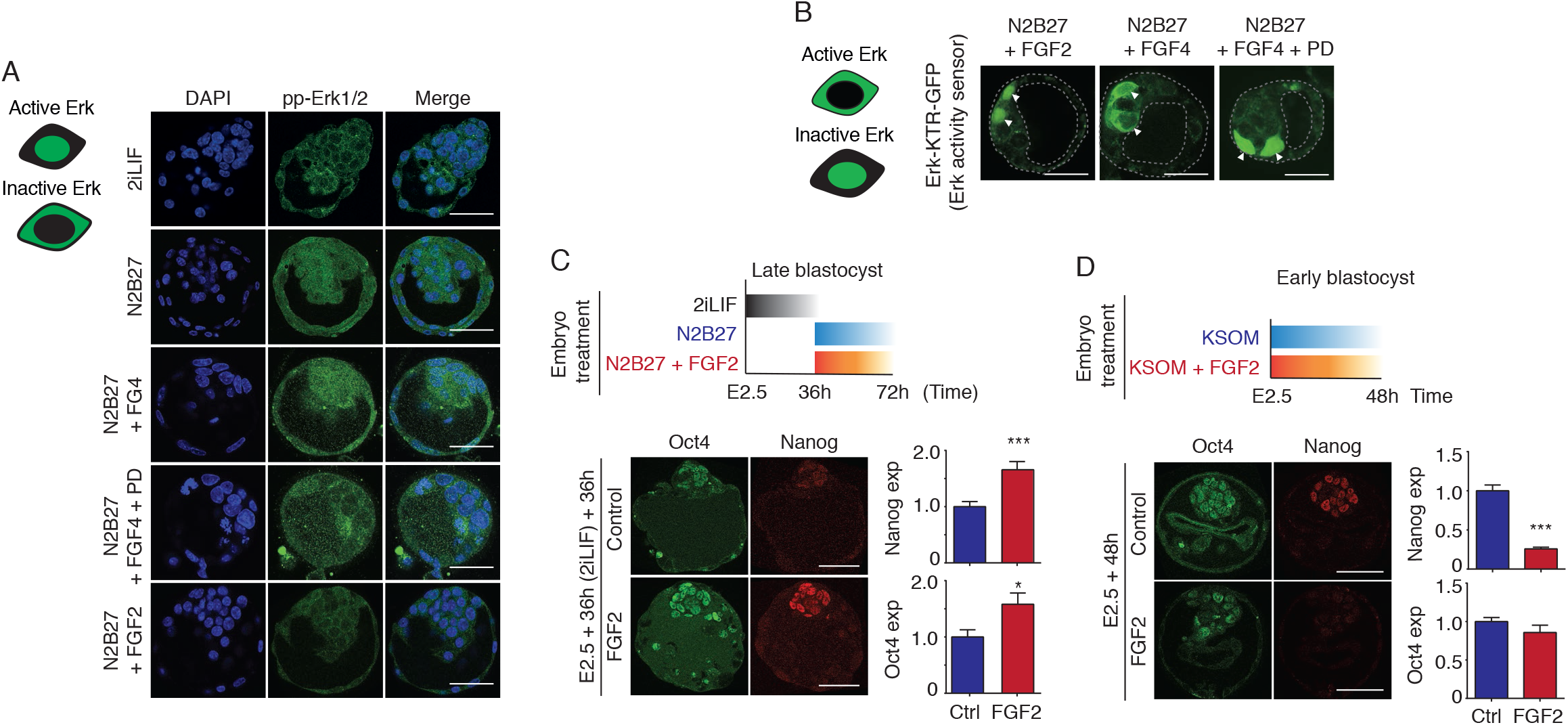
FGF2 inhibits Erk1/2 activity and maintains naïve pluripotency in late blastocysts. (A) Representative images of Erk1/2 activation (pp-Erk1/2) in E4.5 blastocysts treated with N2B27 in the presence or absence of FGF2 or FGF4 for 30min. Mek inhibitor PD0325901 (PD) was added at 5μM. Scale bar represents 50μm. n>10 embryos per each condition. (B) Representative images of Erk1/2 activation in E4.5 blastocysts expressing the Erk activity sensor Erk-KTR-GFP and treated with N2B27 in the presence or absence of FGF2 or FGF4 for 30min. Mek inhibitor PD0325901 (PD) was added at 5μM. Scale bar represents 50μm. Arrows highlight cells expressing the Erk activity sensor. n>10 embryos per each condition. (C) Representative images and quantification of Nanog and Oct4 expression in the ICM of E2.5 blastocysts cultured to E4 stage in 2iLIF media and treated with N2B27 in the presence or absence of FGF2 for 36h. Scale bar represents 50μm. Error bars show mean ±SD. n>11 embryos per each condition. *** for p< 0.0001. (D) Representative images and quantification of Nanog and Oct4 expression in the ICM of E2.5 blastocysts cultured with KOSM in the presence or absence of FGF2 for 48h. Scale bar represents 50μm. Error bars show mean ±SD. n>10 embryos per each condition. p< 0.0001.

These observations were further confirmed by using the Erk-KTR activity biosensor (Regot *et al*., 2014). Mouse embryos at E2.5 were infected with lentiviral particles containing Erk-KTR-GFP sensor and let develop until E4.5 embryonic stage by *ex-vivo* culture in 2iLIF media. As seen with the ppErk antibody, treating embryos with N2B27 in the presence of FGF4 for 30 minutes caused a significant increase in Erk1/2 activation, as seen by the cytoplasmic localization of the RTK-GFP sensor (Figure 6B). The RTK-GFP sensor became nuclear (i.e. Erk became inactive) if embryos were treated with FGF4 in the presence of a Mek1/2 inhibitor (PD) (Figure 6B). Importantly, treating embryos with FGF2 for 30 minutes resulted in Erk inactivation (i.e. nuclear localization of the sensor) (Figure 6B), strongly supporting a similar role of FGF2 in mES cells and late blastocysts.

We therefore investigated whether FGF2 could promote naïve pluripotency in the epiblast of pre-implantation embryos (i.e. in late blastocysts), where Nanog expression is key to maintain naïve pluripotency. We isolated mouse embryos at E2.5 and developed them into E4 embryonic stage by treatment with 2iLIF media for 36 hours, after which, E4 blastocysts were transferred to N2B27 media with or without FGF2 for further 36h (Figure 6C). Notably, we observed that embryos cultured in presence of FGF2 display a higher level of Nanog and Oct4 comparatively to those treated with N2B27, where levels of Nanog and Oct4 expression were significantly lower (Figure 6C).

We next tested whether this was true for early blastocysts. Previous findings described that FGF signalling in early E2.5 embryos promoted loss of Nanog and up-regulation of PrE-lineage markers during Epi/PrE lineage specification (Kang *et al*., 2013). We therefore subjected E2.5 embryos to KSOM treatment in the presence or absence of FGF2 for 48 hours as previously described and assessed the expression of pluripotency (Nanog and Oct4) and PrE lineage (Gata4 and Gata6) markers (Figure 6D and S6B). Similarly to Kang and colleagues (Kang *et al*., 2013), we saw that in early blastocysts, FGF2 indeed repressed the expression of Nanog, but not of Oct4 (Figure 6D) and led to up-regulation of Gata4 and Gata6 (Figure S6B).

Altogether, these results confirm our observations in mES cells and suggest that FGF2 has the potential to maintaining pluripotency is a time-specific manner, likely restricted to the late blastocyst stage in early embryos.

### FGF2 is expressed in the niche of the pre-implantation embryo

The fact that FGF2 helps maintain naïve pluripotency in late blastocysts presumes that FGF2 should, in principle, be expressed in the epiblast. However, FGF2 is known not to be expressed in the epiblast in pre-implantation blastocysts. This posed a conundrum: if FGF2 is indeed important for maintaining naïve pluripotency what is the source of endogenous FGF2 in late blastocysts?

We hypothesized that perhaps FGF2 could be part of the embryo’s niche, being expressed either in the extraembryonic tissues or expressed maternally thereby signalling to the epiblast from the surrounding tissues.

Maternal expression of LIF, another crucial factor for naïve pluripotency is in fact well documented {Stewart, 1992 #190} and FGF2 has also been postulated to be expressed both in trophoblast cells and in maternal (endometrial) tissues (Wordinger *et al*., 1992; Carlone and Rider, 1993; Taniguchi *et al*., 1998; Paria *et al*., 2001; Yang *et al*., 2015).

We tested this hypothesis first by using trophoblast-derived cell lines (Figure S7A and S7B). We saw that FGF2 is indeed expressed at the protein level in two different trophoblast cell lines where it co-localizes with Cdx2, a well establish trophoblast marker (Figure S7A). In addition, exposing mES cells to trophoblast conditioned media for 24h significantly increased Nanog levels when compared to mES cells growing in either (unconditioned) N2B27 or 2iLIF (Figure S7B).

Importantly, we see that FGF2 is highly expressed at the mRNA level *in vivo*, in the mouse uterus (Figure 7A). Strikingly, we see FGF2 levels enriched in the uterus of day E4.5 pregnant mice, comparatively to other tissues, such as, liver or thymus or in the uterus of non-pregnant mice (Figure 7B).

**Figure 7.**
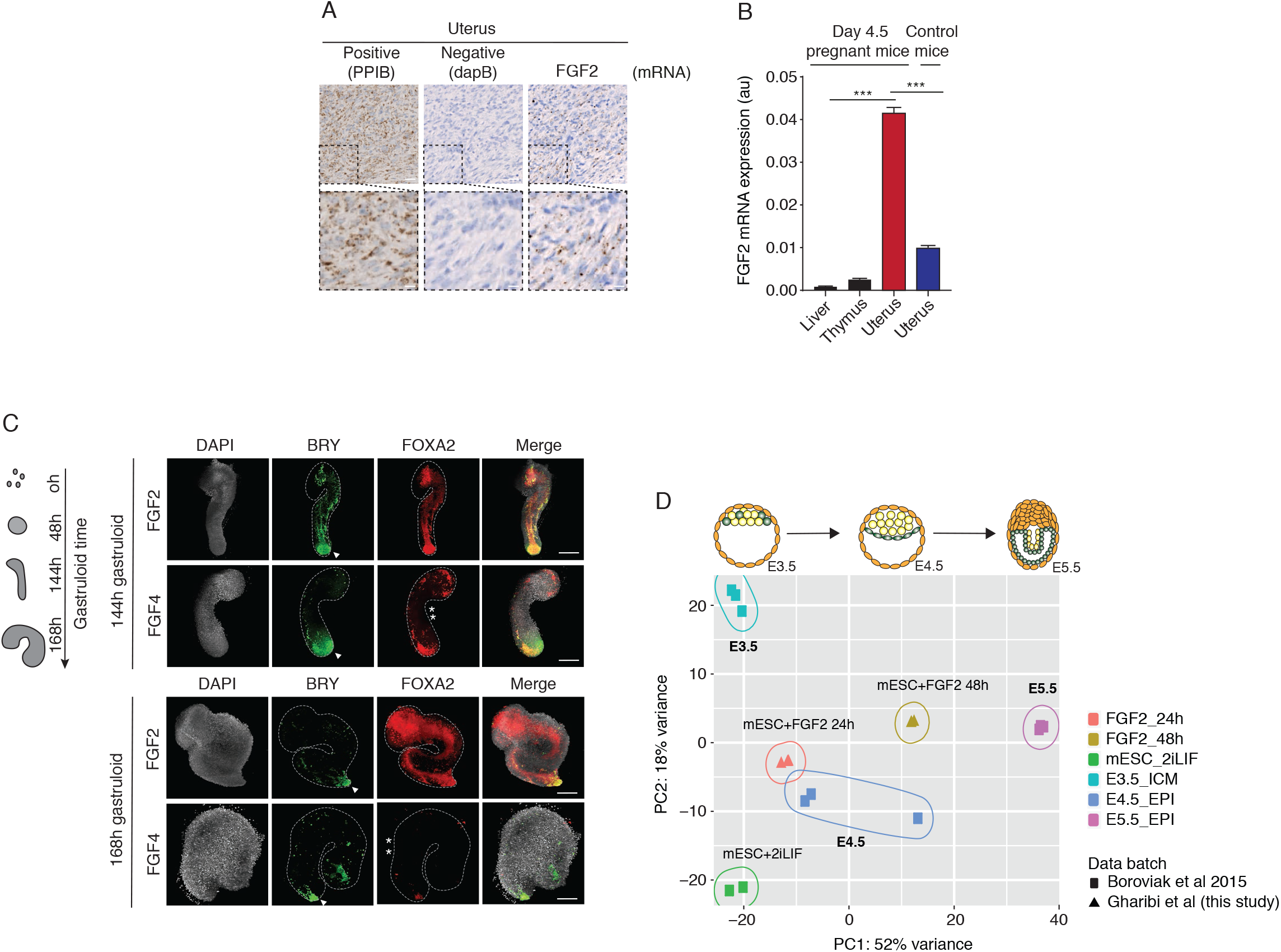
FGF2 is expressed in the niche of the pre-implantation embryo and may be important for coordinating optimal lineage proportions during cellular differentiation. (A) Relative expression levels of FGF2 in the liver, thymus and uterus tissues of E4.5 pregnant mice. Uterus from non-pregnant mice was used as control (blue). Data was normalized to house-keeping gene GAPDH. n=3 independent experiments. The error bars show mean ±SD. (B) Representative RNAscope images of mRNA staining in a mouse uterus showing PPIB as positive control, dapB as negative control and FGF2. Probe binding is visualised as punctate brown dots. Scale bar represents 50μm and 10μm (inset). (C) Top: Representative images of mouse gastruloids at 144h resulting from mES cells treated with FGF2 (100ng/ml) or with FGF4 (100ng/ml) for the first 48h. Primitive streak marker, Brachyury (BRY) is shown in green and endoderm marker, FOXA2, is shown in red. Arrows show BRY localization at the pole. Stars highlight lack of FOXA2 tubular staining. Scale bar represents 100μm. Bottom: Representative images of mouse gastruloids at 168h resulting from mES cells treated with FGF2 (100ng/ml) or with FGF4 (100ng/ml) for the first 48h. Primitive streak marker, Brachyury (BRY) is shown in green and endoderm marker, FOXA2, is shown in red. Arrows show BRY localization at the pole. Stars highlight lack of FOXA2 tubular staining. Scale bar represents 100μm. (D) Principle component analysis plot based on differentially expressed genes (LRT) after batch effect removal, comparing in vivo mRNAseq data from E3.5 (ICM), E4.5 (EPI) and E5.5 (EPI) blastocysts and mES cells treated with 2iLIF from Boroviak et al 2015 and mRNAseq data from FGF2 treated mES cells for 24 and 48h, from this study.

Together these results show that FGF2 is expressed maternally and in extra-embryonic tissues and suggests it is part of the niche of the early pre-implantation embryo.

### FGF2 may contribute to generating lineages with the right cell number during early mouse development

Finally, we wanted to understand how FGF2 signals from the niche may impact lineage specification in early embryogenesis.

In order to address this, we generated mouse FGF2 and FGF4-treated gastruloids, a new 3D in-vitro system that recapitulates early mouse development (Beccari *et al*., 2018, 2019). Both FGF2 and FGF4 treated mES cells gave rise to gastruloids that survived at least 7 days (Figure 7C). Imaging FGF2 and FGF4-treated gastruloids by light-sheet microscopy showed that these gastruloids have similar shapes and undergo extremely similar development, including the formation of the primitive streak, one of the first signs of gastrulation, marked by the expression of Brachyury (BRY) (Figure 7C).

However, we saw that while we could observe a robust tubular expression of FoxA2, a canonical marker of embryonic endoderm in FGF2-treated gastruloids, FGF4-treated gastruloids showed a markedly reduced expression of FoxA2 at 144h (Figure 7C). This is unlikely to be just a delay in gastruloid development because examining later gastruloids at 168h, in which Brachyury expression has decreased for both FGF2 and FGF4 treatments, showed an even more remarkable difference in FoxA2 expression between the two treatments.

This striking observation suggests that FGF2 may help coordinate cell number and optimal proportion of specific embryonic lineages during early embryonic development.

### FGF2 likely inhibits Erk1/2 in *in-vivo* thereby promoting naïve pluripotency

The remaining question was thus whether FGF2 could be the factor that inhibits Erk1/2 in-vivo, helps keep Nanog and pluripotency factors up-regulated and maintain a signature of low DNA methylation, both important hallmarks of naïve pluripotency in vivo.

In order to address this, we performed a global RNA sequencing (RNAseq) analysis of mES cells treated with FGF2 for 24 and 48 hours, and compared the resulting genomic signatures with published RNAseq in vivo data from early mouse embryos (Boroviak *et al*., 2015) (Figure 7D). We saw that developmental time of the early embryo is well captured in principal component number 1 (PC1), where most variance in gene expression data originates from. As expected, mES cells treated with 2iLIF have a gene expression signature falls close to E4.5 embryos. However, we see that FGF2-treated cells for 24h are remarkedly more similar to E4.5 embryos (Figure 7D). Interestingly, longer treatment of mES cells with FGF2 for 48h results in gene expression patterns that lie between embryonic days E4.5 and E5.5, mirroring transient gene expression seen during this embryonic transition *in vivo* (Figure 7D).

This strongly suggests that FGF2-treated mES cells resemble the transient naive pluripotency cells in the epiblast of late blastocysts.

Altogether, our findings support that endogenous FGF2 in surrounding tissues of the pre-implantation embryo may help support naïve pluripotency and that FGF2 may be the factor that inhibits Erk1/2 activity in vivo, thereby helping support a transient transition through pluripotency states and potentially optimal lineage specification in the early embryo.

## Discussion

Embryonic development is regulated by precise temporal integration of signalling cues that promote ordering of events and commitment to lineage specification. As such, FGF signalling has been widely believed to be an important pro-differentiation signal by activating Erk1/2, and hence detrimental to the maintenance of pluripotency.

The evidence presented here argues that FGF2 is instead be a key factor maintaining low Erk1/2 activity and consequently promoting naïve pluripotency in mES cells and late blastocysts.

We found that despite being an Erk1/2 activator, and in sharp contrast to other FGF family members, such as FGF4, FGF2 only transiently activates Erk1/2 and induces instead, a persistent inhibition of Erk1/2 activity (Figure 1).

Since the establishment of the first mES cell lines a few factors have been described to be at the heart of maintaining naïve pluripotency, including inhibition of Erk1/2 activity. However how Erk1/2 is inactivated in vivo has remained a mystery.

We propose that the molecular mechanism by which FGF2 induces Erk1/2 inactivation relies on an incoherent feedforward regulation whereby FGF2 brings about both the activation of Erk1/2 and the inhibition of Ret, a potent Erk1/2 activator (Figure 2). Ret is highly expressed in early embryogenesis, starting at the eight-cell stage until peaking at the blastocyst stage (Li *et al*., 2009). We see that simultaneous activation of both Ret and FGF pathways leads to a full (sustained) Erk1/2 activation and promotes pluripotency exit. This mechanism was further validated by finding that FGF4 in the presence of Ret inhibitors successfully mimicked the effect of FGF2 on Erk1/2 activity (Figure 2). In line with these observations, a recent study showed that a double knock-out of Fgfr1^-/-^; Fgfr2^-/-^ failed to completely block differentiation of ES cells, indicating that other pathways are able to compensate for the lack of FGF signalling (Molotkov *et al*., 2017).

We propose that FGF2 down-regulation of Ret activity is likely to be driven by endocytosis, which explains why simultaneous stimulation of cells with FGF2 and FGF4 results in a transient Erk1/2 activation (Figure 2). Receptor tyrosine kinase co-regulation is not specific to embryonic development but a recurrent phenomenon in signal transduction (Stallaert et al., 2018; Latko et al., 2019).

FGF2 has been implicated in the self-renewal of human embryonic stem (hES) cells and mouse epiblast stem cells (EpiSCs) (Greber *et al*., 2010) and shown to increase the efficiency of EG cell derivation, where it maintains long-term expression of pluripotency markers. In addition, and consistent with the idea that FGF2 supports pluripotency, recent evidence has revealed that reprogramming efficiency could be improved by FGF2 (Chen *et al*., 2010; Han *et al*., 2011; Jiao *et al*., 2013).

We see that FGF2-induces up-regulation of naive pluripotent factors in mES cells (Figure 3) and down-regulation of DNA methylation (Figure 4), both hallmarks of naïve pluripotency in the epiblast (De Los Angeles *et al*., 2015). Importantly, FGF2 treated mES cells were able to contribute to the generation of chimeras showing that FGF2 maintains pluripotency in an in-vivo context. (Figure 3).

The epiblast is a transient population of cells from which the entire mammalian foetus is derived. mES cells growing in 2iLIF resemble cells of the epiblast and perpetuate a state of naïve pluripotency. However, naïve pluripotency is only a *transient* state *in vivo*, lasting about 24h. Moreover, it has recently been shown that sustained Erk inhibition in 2iLIF conditions leads to *sustained* downregulation of methyltranferases and causes irreversible epigenetic and genomic changes that impair developmental potential (Choi *et al*., 2017). In line with this, our work shows that there is a striking difference between Mek expression levels in 2iLIF and FGF2 treated mES cells (Figure 1E) and that the effect of FGF2 on pluripotency is transient - FGF2 can rescue naïve pluripotency conditions only for 24h, after differentiation initiation. (Figure 5).

We find that the ability of FGF2 to help support naïve pluripotency is likely to be true *in-vivo*. In particular, we see that FGF2 can inhibit Erk1/2 *in vivo* and upregulate naïve pluripotency factors such as Nanog in E4.5 mouse blastocysts (Figure 6). Furthermore, we see that mES cells treated with FGF2 show a very similar global gene expression signatures compared to E4.5 embryos (Figure 7) supporting the idea that FGF2 is likely to be the factor that inhibits Erk *in vivo*.

While FGF2 is known to promote implantation, a non-conditional, homozygous FGF2 deletion gives rise to viable progeny with no apparent pre-implantation phenotype (Ortega *et al*., 1998). This suggests potential compensatory mechanisms and is in line with the fact that maintenance of naïve pluripotency in the embryo is known to dependent on the synergy between multiple factors (i.e. Erk inhibition and activation of Wnt and STAT pathways).

FGF2 is believed not to be expressed in the epiblast and our work shows that FGF2 is expressed both maternally and potentially in the embryo’s supporting tissues. This suggests that FGF2 may be part of the embryo niche (Figure 7). This is supported by several reports (Wordinger *et al*., 1992; Carlone and Rider, 1993; Taniguchi *et al*., 1998; Paria *et al*., 2001; Yang *et al*., 2015). Interestingly, this is reminiscent of the maternal expression of LIF, a key regulator of both naïve pluripotency and implantation (Stewart *et al*., 1992; Ying *et al*., 2008), highlighting that the early embryo may integrate different signals from the its niche.

It is therefore likely that Erk activity may be modulated by different factors during early development. Interestingly, we see that cells exposed to sequential treatment of FGF2 and

FGF4 cues maintain a memory of Erk inhibition, even when FGF4 is 1000x more abundant (Figure S7C). We thus predict that fine-tuning Erk1/2 signalling through internal and external factors might be at the heart of the balance between pluripotency, self-renewal and differentiation during embryonic transitions.

Finally, using an in vitro model of gastrulation we show that exposure to FGF2 affects the number of cells and the proportions of specific lineages, namely endoderm, where Nanog expression takes central place (Figure 7). We predict that signals from the embryo’s niche might contribute to establishing the right proportion of cells during early gastrulation. This beautifully illustrates how a transient signal may give rise to long term fate choice. In line with this, a recent report has shown that FGF signalling couples fate decision to lineage composition and plays a key role in establishing Epi/PrE lineages with the right proportions in early blastocysts (Saiz *et al*., 2020).

In conclusion, we propose a role for FGF signalling inhibiting Erk1/2 and thereby helping support a transient naïve pluripotency state. We further suggest that maintenance of naïve pluripotency and specification of early embryonic lineage might be affected by both internal and external factors, involving signals from the embryo as well as its niche i.e. maternal and extra-embryonic tissues.

Our work highlights how differences in Erk dynamics can impact on fate-choice during early development and how feed-forward regulation can allow for progression through a continuum of transient states, mediating time-dependent integration of signals and thereby facilitating progression through developmental stages. Given how common feedforward regulation is in regulatory networks, we anticipate that coherent and incoherent feed-forward control may prove to be a recurring theme during transient, reversible transitions in early development.

## Authors contributions

BG designed experiments, performed most of the experimental work and data analysis and helped write the paper. EG and PB analysed the proteomics data and AM and HK performed the proteomics experiments. BN, PH, KM and IR helped with mouse embryo experiments. SS and RS helped with data analysis. SDMS conceived the study, helped design experiments and wrote the paper.

## Acknowledgments

We thank the staff at the Flow Cytometry and High-Throughput Screening STPs at the Francis Crick for excellent technical support. We are very grateful to Frank Uhlmann, Matthias Merkenschlager, Peter Sarkies, Peter Hill and the members of the Santos lab for helpful advice and discussions. We are very grateful to Rakee Chauhan for advice and for sharing Ret reagents, Teresa Rayon, Claudia Gerri and Sophie Brumm for advice on mouse embryos, Patty Wai and Jonny Kohl for providing mouse tissues and the Turner and Smith labs at the Crick for sharing the lnx090915.02 mouse trophoblast cell line. We thank James Briscoe and Adrienne Sullivan for critical reading of the manuscript. BN was supported by Inner Mongolia University High-Level Talents award 21400-5165159 and NSFC31760335. The Santos lab at the Francis Crick Institute is supported by funding from Cancer Research UK (FC001596), the UK Medical Research Council (FC001596) and the Welcome Trust (FC001596).

---

**Supplemental Figure S1.**
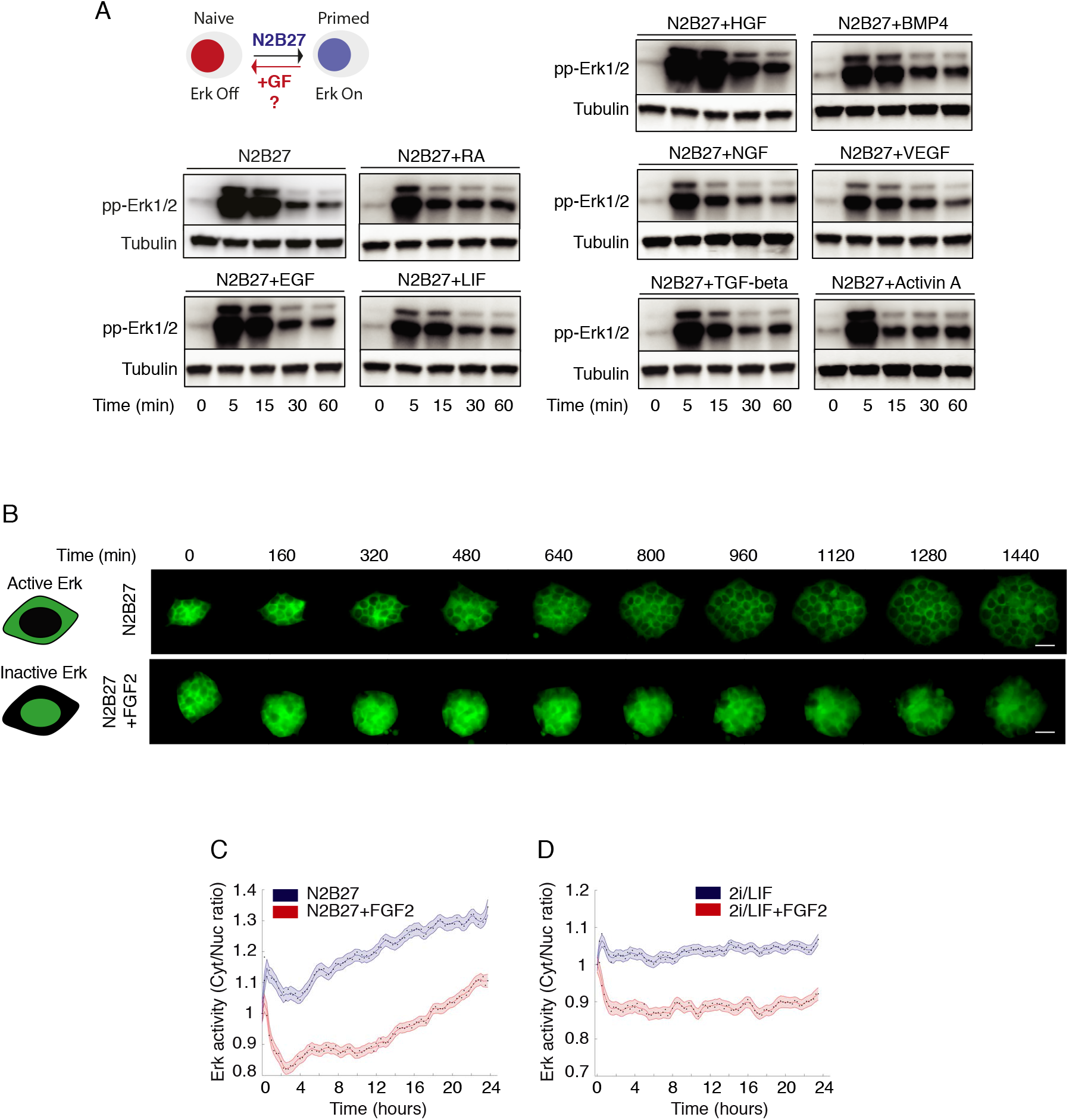
FGF2 induces transient activation and long-term inhibition of Erk1/2 in mES cells (related to Figure 1) (A) Western blot time courses comparing Erk1/2 (pp-Erk1/2) activation following treatment of N2B27 with the indicated growth factors (GF). (B) Schematic of the Erk-KTR activity sensor used to measure Erk1/2 activity in single live cells and representative images of a long, 24h time course of mESCs treated with N2B27 in the presence or absence of FGF2. Scale bar represents 20μm. (C) Erk activity over 24h as measured in single Erk-KTR mESCs in N2B27 in the presence or absence of FGF2. C/N ratio indicates ratio of cytoplasmic to nuclear intensities. n > 100 cells were analyzed for each experimental condition. (D) Erk activity over 24h as measured in single Erk-KTR mESCs in 2iLIF in the presence or absence of FGF2 (100ng/ml). C/N ratio indicates ratio of cytoplasmic to nuclear intensities. n > 100 cells were analyzed for each experimental condition.

**Supplemental Figure S2.**
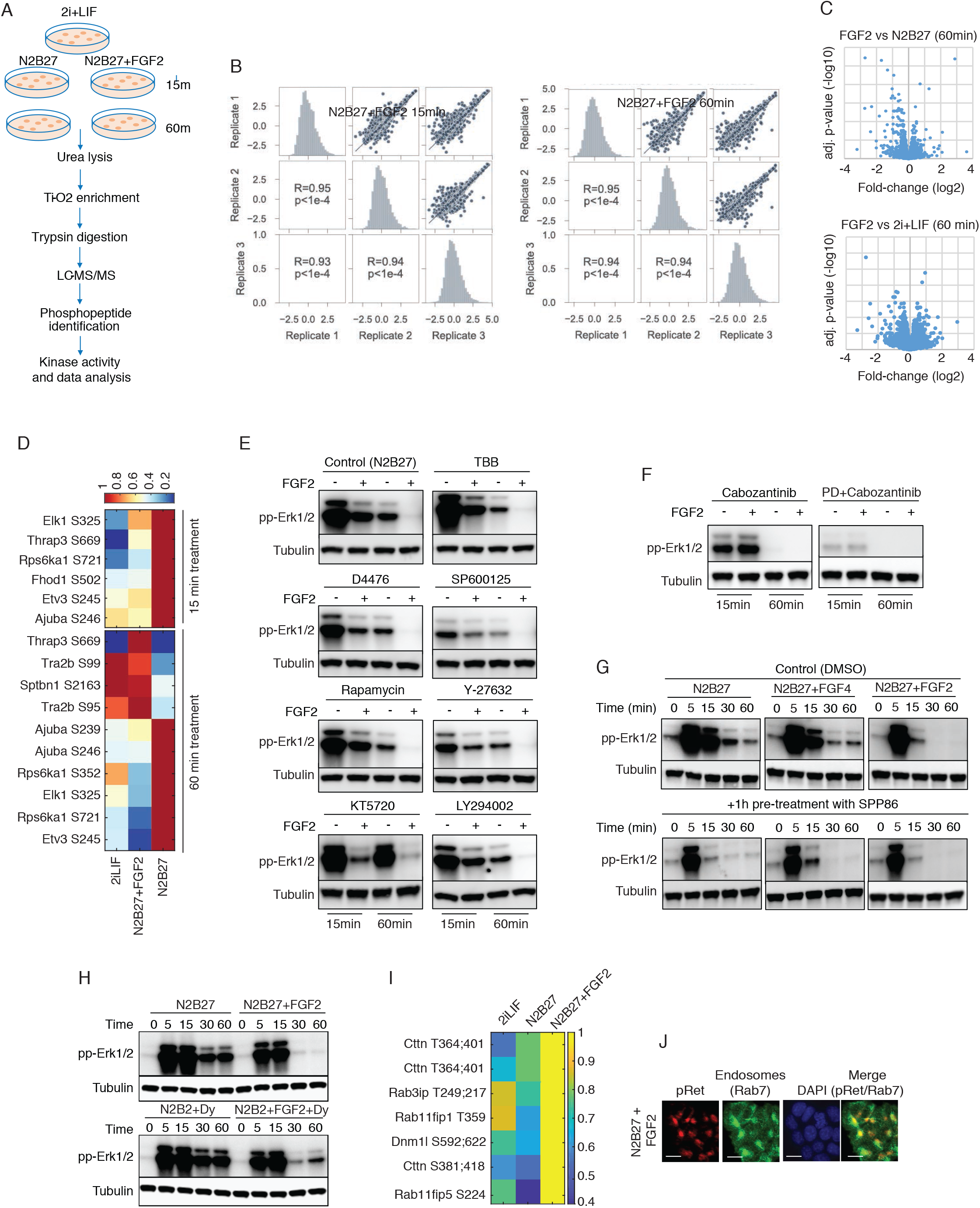
FGF2 induces a transient Erk1/2 activation via a Ret-dependent incoherent feedforward loop (related to Figure 2) (A) Schematic of the phosphoproteomics experimental work flow. (B) Correlation analysis for the three biological replicates for two different time points of the N2B27+FGF2 treated samples analyzed by phosphoproteomics. (C) Volcano plots comparing adjusted p-value (-Log10) as a function of the log2 changes in phosphopeptide abundance in 2iLIF or N2B27 (control) and N2B27+FGF2 (FGF2) treated samples for 15 and 60min. (D) Heatmaps representing changes in phosphorylation of Erk1/2 targets in cells treated with either 2i LIF, FGF2 or N2B27 for 15 and 60 minutes. (E) Western blot analysis of Erk 1/2 activation following treatment with inhibitors of kinases predicted by KSEA analysis. Representative of two independent experiments. (F) Western blot analysis of Erk 1/2 activation following Ret inhibition using Cabozantinib in the presence or absence of FGF2 (left) or following Ret and FRFR inhibition using Cabozantinib and PD173074, respectively, in the presence or absence of FGF2. Representative of two independent experiments. (G) Western blot time of Erk 1/2 activity over time following stimulation with N2B27 alone or in presence of FGF4 or FGF2 in cells pre-treated with Ret inhibitors SPP86 or Vandetanib. n = 2 independent experiments. (H) Western blot of Erk 1/2 activity over time following stimulation with N2B27 alone or N2B27+FGF2 in presence or absence of endocytosis inhibitor dynasore (DY). n = 2 independent experiments. (I) Heatmaps representing changes in phosphorylation of endocytosis regulators in cells treated with either 2i LIF, N2B27 or N2B27+FGF2 for 60 minutes. (J) Representative immune-fluorescence images of active Ret (pRet) and Rab7 (early endosomes marker) colocalization after FGF2 stimulation. Scale bar represents 10μm.

**Supplemental Figure S3.**
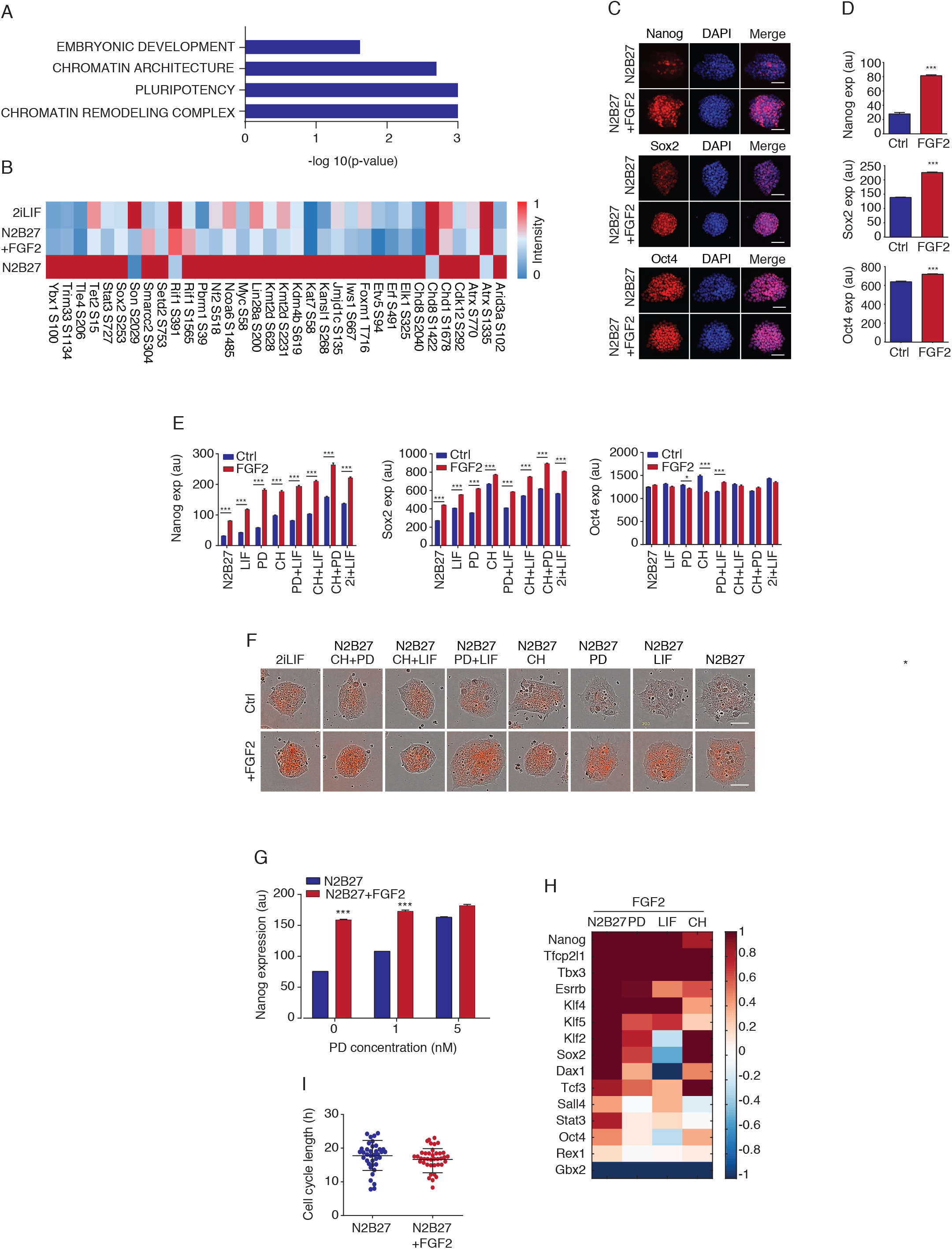
FGF2 supports naïve pluripotency in mouse ES cells. (A) Subset of the gene ontology (GO) terms enrichment analysis for the differently regulated phosphopeptides in FGF2 treated samples. (B) Heatmap comparing changes in phosphorylation of pluripotency regulators after 2i LIF, N2B27 or N2B27+FGF2 treatments from differentially expressed phosphopeptides. Colour scale shows minimum and maximum signal intensity as 0 and 1, respectively. (C) Representative immunofluorescence images of core pluripotency factors in E14 ES single cells cultured for 48h in N2B27 with or without FGF2. Scale bar represents 50μm. (D) Quantification of core pluripotency factors in E14 ES single cells cultured for 48h in N2B27 with or without FGF2. n> 500 cells were analysed for each experimental condition. The error bars show mean ±SEM. *** p<0.0001 using KS- and Mann-Whitney tests. (E) Analysis of core pluripotency factors in cells treated with N2B27 alone or with different combination of 2i LIF components in presence or absence of FGF2 (100ng/ml) for 48h. n>500 cells were analysed for each experimental condition. The error bars show mean ±SEM. *** p<0.0001. This is representative of n > 3 independent experiments. (F) Representative images of Nanog-mCherry stable mES cells cultured for 24h under different combinations of 2i LIF components in presence or absence of FGF2. Scale bar represents 50μm. (G) Nanog expression as a function of Erk inhibitor PD concentration (PD) in the presence or absence of FGF2 (100ng/ml). n>500 cells were analysed for each experimental condition. The error bars show mean ±SEM. *** p<0.0001. (H) Gene expression of pluripotency associated transcription factors in ES cells cultured in N2B27 with either PD (1 μM), CH (3 μM) or LIF (10ng/ml) in the presence of FGF2 for 48h. Data was normalized to GAPDH and shown as log2 fold change relative to the corresponding treatment in the absence of FGF2, which was set to 0. n = 3 independent experiments. (I) Cell cycle duration as measured by two consecutive mitoses in cell cultured in N2B27 in the presence or absence of FGF2. n > 40 cells were analysed for each experimental condition.

**Supplemental Figure S4.**
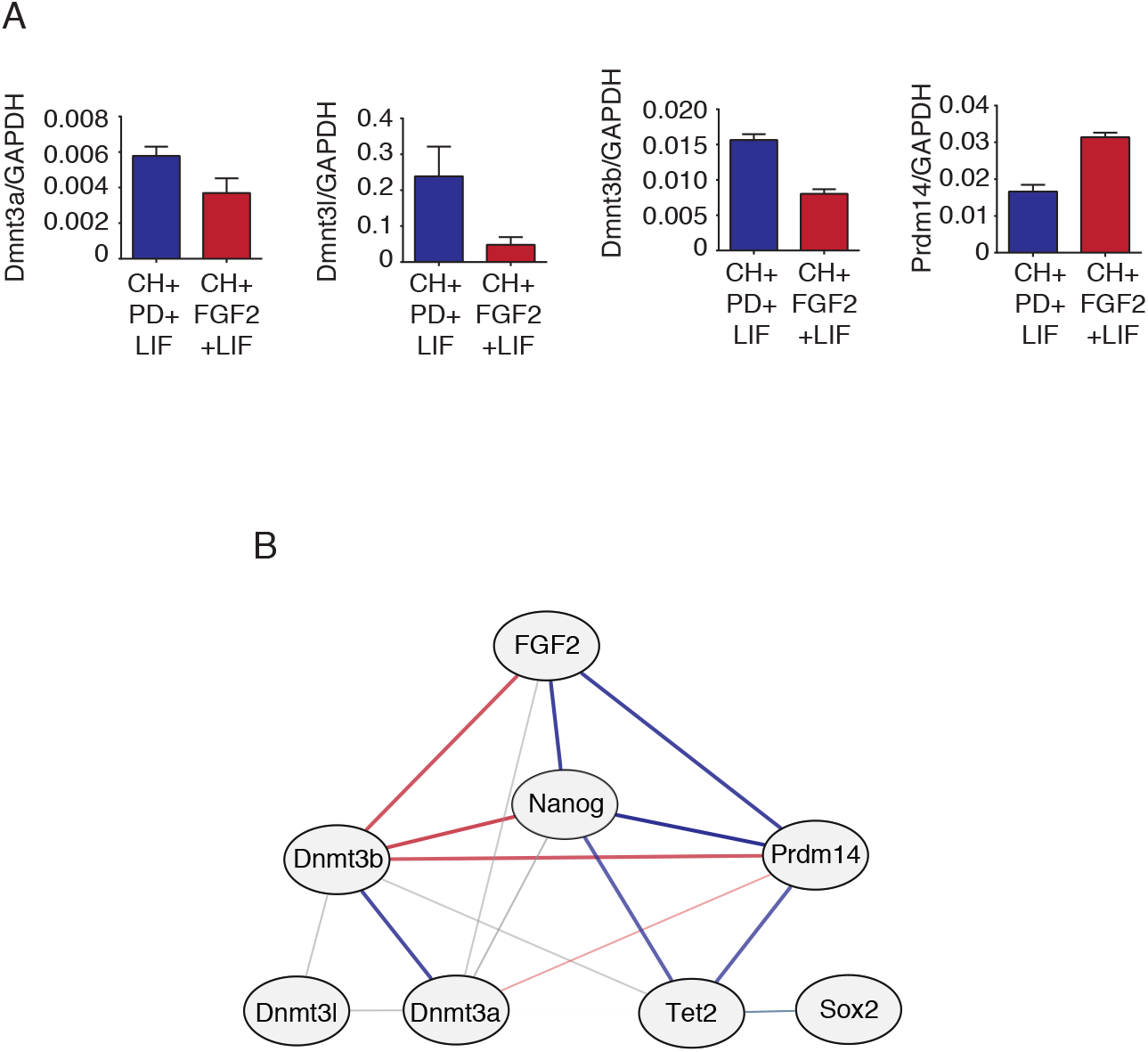
FGF2 supports naïve pluripotency in mouse ES cells by regulating DNA methylation (related to Figure 4) (A) Expression of DNA methylation associated genes by qPCR after culturing ES cells in N2B27 with CH and LIF in presence of PD or FGF2 for 5 passages. n = 2 independent experiments. The error bars show mean ±SD. (B) Computed network of possible interactions between FGF2, pluripotency and epigenetic genes. Positive interaction is represented by blue line; negative interaction is indicated by red line; the thickness of each line reflects the degree of regulation.

**Figure S5.**
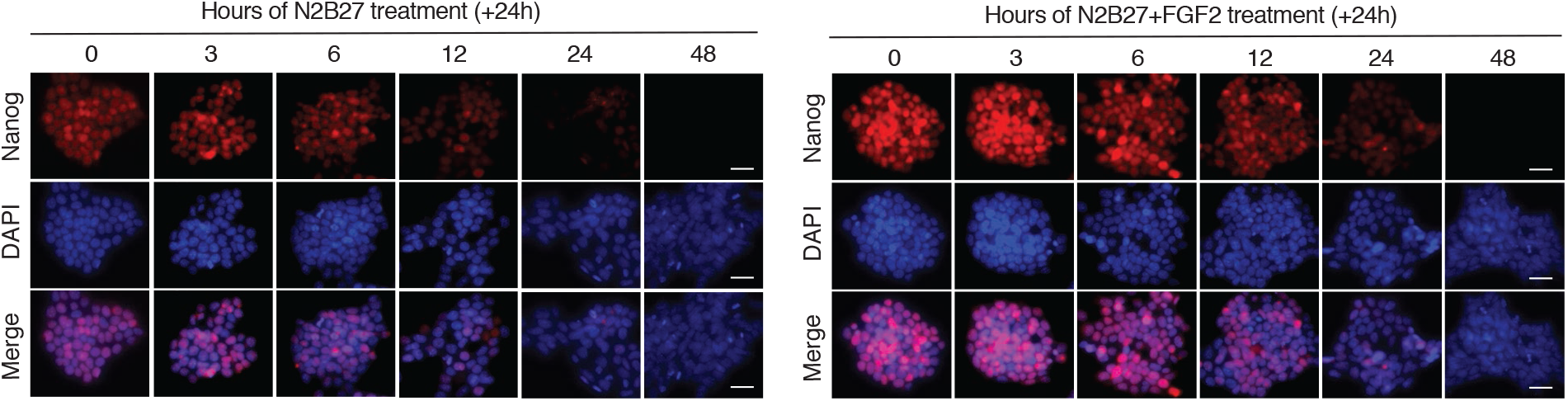
FGF2 rescues and sustains Nanog expression for up to 24h following N2B27 treatment (related to Figure 5) Representative images of Nanog expression in cells primed in N2B27 for indicated time and subsequently treated for 24h in N2B27 in the presence or absence of FGF2. Scale bar represents 20μm.

**Figure S6.**
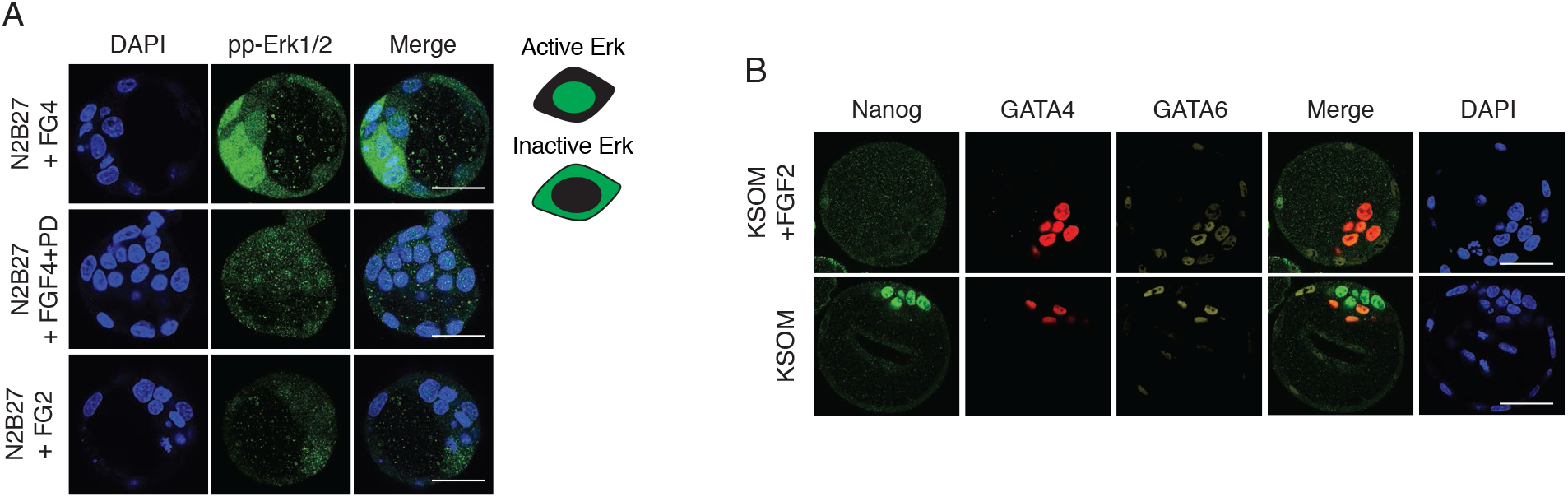
FGF2 maintains Erk1/2 inactive in E4.5 stage blastocysts (related to figure 6) (A) Erk activity in FGF4 and FGF2 treated E4.5 blastocysts using antigen retrieval IF technique of a phospho-specific Erk1/2 antibody. FGF4 in the presence of PD0325901 (5μM) was used as a control. Scale bar represents 50μm. n>10 embryos per each condition. (B) Representative images of Epi (Nanog) and PrE (Gata6 and Gata4) markers expression in E2.5 embryos cultured in the presence or absence of FGF2 for 48h. Scale bar represents 50μm.

**Supplemental Figure S7.**
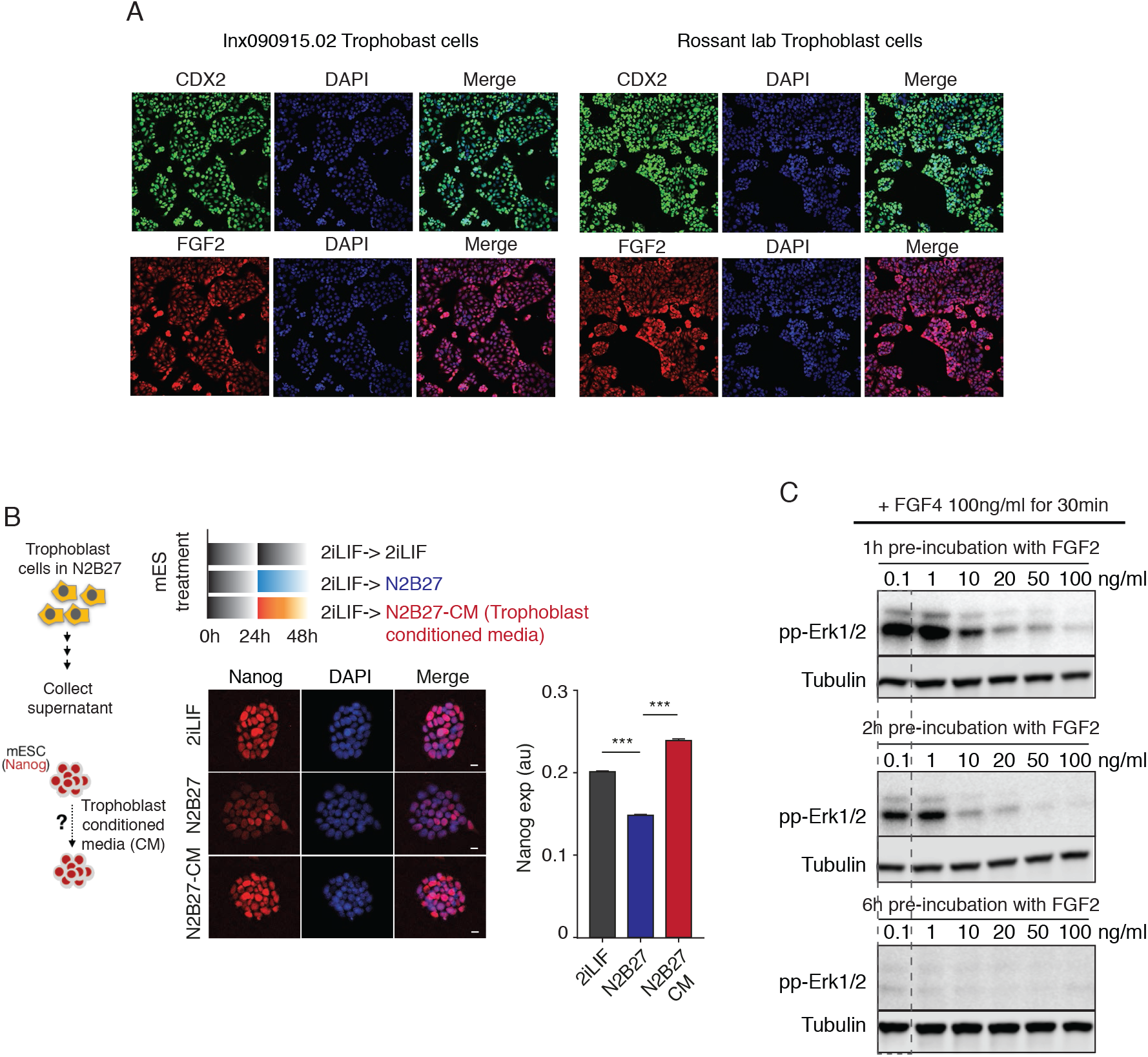
FGF2 is expressed in the niche of the pre-implantation embryo and may be important for sustaining a diapause signature of c-Myc and Ki67 inhibition (related to figure 7 and discussion) (A) Representative images of co-expression of the canonical trophoblast marker CDX2 and FGF2 in two trophoblast cell lines: Lnx090915.02 trophoblast cells (left) and trophoblast cells obtained from the Rossant lab (right). Scale bar represents 50μm. (B) Representative images and quantification of endogenous Nanog expression in R1-Nanog mES cells cultured for 24h in 2iLIF, N2B27 or N2B27 conditioned media from trophoblast stem cells (TS). Scale bar represents 20μm. n>500 cells were analyzed for each experimental condition. p< 0.0001. The error bars show mean ±SEM. (C) Western blot time courses comparing Erk1/2 activity (pp-Erk1/2) in mES cells treated with FGF4 (100ng/ml) for 30min after pre-treatment of cells with FGF2 (100ng/ml) for increasing lengths of time: 1h (top), 2h (middle) and 6h (bottom). α-Tubulin was used as loading control for quantification. Representative of n=2 independent experiments.

