## Supplemental_Information for "A FGF2-mediated incoherent feedforward loop induces Erk inhibition and promotes naïve pluripotency"

Gharibi et al 2020

### Materials and Methods

#### Growth factors, inhibitors and antibodies

Growth factors used in this study were: FGF-basic recombinant mouse protein (1-100 ng/ml), FGF-basic (AA 10-155) recombinant human protein (1-250 ng/ml), FGF4 recombinant human protein (100 ng/ml), BMP4 (100 ng/ml), VEGF and TGF-B (100 ng/ml) (Thermo Fisher scientific), FGF5 (100 ng/ml) (Abcam), EGF (100 ng/ml), NGF (100 ng/ml) and HGF (100 ng/ml) (PeproTech), FGF8 (100 ng/ml) and LIF recombinant mouse protein (100 ng/ml) (Biolegend), FGF3 (100 ng/ml) (Medsystems Ltd), Retinoic Acid (5μM) (Sigma), Activin A (100 ng/ml) (Cambridge Biosciences).

Inhibitors used in this study were: PD0325901 (1-10 μM) and vandetanib (10 μM) (Cayman Chemical), Dynasore (100 μM) and CHIR99021 (3 μM) were from Sigma, wortmannin (1 μM) (Calbiochem) D4476 (20 μM) (BioVision), Y-27632 (10 μM) (VWR), SPP 86 (10 μM), Go 6983 (5 μM), TBB (10 μM) (Tocris), Rapamycin (10 μM), 10058-F410 (50μM) (Thermo Fisher scientific), KT5720 (1 μM) (Generon), SB203580 (20 μM), SB-431542 (10 μM), PD173074 (10 μM), SP600125 (10 μM) and LY294002 (10 μM) from Cell Guidance Systems and PKC CAMK inhibitor cocktail (1:500) from Millipore.

**Table 1.** Antibodies used in this study

| Primary antibodies | Source | Catalog nr/RRDI | Dilution |
|---|---|---|---|
| 5-mC antibody | CST | 28692/ AB_2798962 | 1:1500 |
| DNMT3B antibody | Novus | NB100-56514/ AB_838139 | 1:200 |
| c-MYC | CST | 5605/AB_1903938 | 1:100 |
| GATA6 antibody | R&D | AF1700 | 10ug/ml |
| Ki-67 antibody | R&D | 50-5698-82 | 10ug/ml |
| Nanog antibody | Abcam | ab80892 | 1:200 |
| Nanog antibody | R&D | 14-5761-80 | 2ug/ml |
| Oct4 antibody | Abcam | ab19857 | 1:500 |
| phospho-Akt (Ser473) | CST | 4060 | 1:1000 |
| phospho-C-Raf (S338 | CST | 9427 | 1:1000 |
| phospho-JNK1+JNK2 (T183 + Y185) | Abcam | ab4821 | 1:1000 |

| | | | |
|---|---|---|---|
| phospho-p38 MAPK (Thr180/Tyr182) | CST | 4511 | 1:1000 |
| phospho-MAPK p42/p44 (T202/Y204) | CST | 4370 | 1:2000 |
| phospho-PKC (S660) | CST | 9371 | 1:1000 |
| phospho-RSK-1/2 (Thr359/Ser363) | Santa Cruz | sc-12898-R | 1:500 |
| Sox2 antibody | Abcam | ab97959 | 1:500 |
| α-tubulin | Sigma | T5168 | 1:4000 |
| Secondary antibodies | Source | Catalog nr | Dilution |
| Alexa 488 goat anti-rabbit IgG | TFS | A11008 | 1:200 |
| Alexa 594 goat anti-mouse IgG | TFS | A21203 | 1:200 |

**Cell culture**

R1, E14 mouse embryonic stem cells (mESC) were cultured without feeders on gelatine (0.1%) coated dishes and routinely maintained in serum free N2B27 media (1:1 ratio of Neurobasal: DMEM/F-12, N2 (1:200), B27 (1:100), penicillin/streptomycin, 2 mM L-glutamine and 0.05% BSA) supplemented with 3 μM CHIR99021, 1 μM PD0325901 and 10ng/ml Leukemia Inhibitory Factor (LIF). For particular experiments as stated in the main text cell were grown in serum containing media consisting of Knockout™ DMEM supplemented with 10% FCS, 1 × MEM non-essential amino acids, penicillin/streptomycin, L-glutamine, 0.05 mM β-mercaptoethanol and 10ng/ml LIF.

**Biosensors and generation of stable cell lines**

The Nanog-mCherry knock-in reporter mES cell line was generated by CRISPR/Cas9 system as previously described (Yang et al., 2013). The bicistronic expression vector pX330 (pX330, Addgene plasmid # 42230) was used to express Cas9 and a sgRNA. Briefly, sgRNA (5′- CACCGCGTAAGTCTCATATTTCACC-3′ and 5′-AAACGGTGAAATATGAGACTTACGC-3′) were cloned at BbsI site into the px330 and co-transfected along with previously described. Nanog-2A-mCherry donor vector in to mouse R1 ES cells using TransIT-LT1 Transfection Reagent according to manufacturer instruction. Cells were expanded and sorted for mCherry positive population.

The ERK KTR-GFP fragment was PCR amplified from previously described pLentiCMV Puro DEST ERKKTRClove (Regot et al., 2014) and cloned into lentiviral vector CSII-EF-1-MCS-2 by restriction digestion and ligation reactions. The CSII-EF-1-MCS-2 plasmid is a modified CSII-EF-1-MCS backbone vector where the linker TCGAAGCTAGCCCTGCAGGTTAATTAAC has been added to the MCS to increase the number of unique restriction sites.

The B6N6-H2B-mCherry stable mES cell line was generated by lentiviral infection of pcs-EF1α-H2B-mCherry. Lentivirus production was carried out in 293T cells transfected with lentiviral vector and lentivirus assembly vectors (PAX2 and VSVG) using with Polyethylenimine (PEI). Packaged lentivirus were harvested after 48h and used to infect cells in presence of polybrene. Cells were expanded and sorted on a Becton Dickinson FACSAria III influx to obtain pure populations expressing the desired fluorescent reporters.

**shRNA knockdown**

shRNA knockdown was carried out using GIPZ lentiviral shRNA (Thermo Scientific). shRNA sequences were: Nanog, AGCCAGGTGGGCAAAGAACTA, SOX2 and negative control for nonspecific effects (NC) ggaatctcattcgatgcatac (catalog no. 336313KH1172N). The lentiviral particles were produced in 293T using lentivirus assembly vectors (PAX2 and VSVG) and Polyethylenimine (PEI). Packaged lentivirus were used to infect cells in presence of polybrene. Cells were cultured for 48h and selected with 1μg/ml of puromycin.

Nanog and Sox2 shRNA lines were used to compute gene co-expression network in N2B27 and N2B27+FGF2 conditions. Under these perturbations, the levels of Nanog, Dnmt3a, Dnmt3b and Dnmt3l, Tet2, Sox2 and Prdm14 were quantified using qRT-PCR. Pearson correlation co-efficient was calculated between the expression of all proteins-pairs and was then used to construct a Pearson co-expression network (as described in (Dunn et al., 2014)).

**Mouse embryo experiments and generation of chimeras**

All animal were procedures including breeding and maintenance were performed under a UK Home Office Project License and according to the UK Home Office regulations and United Kingdom Animals (Scientific Procedures) Act (1986).

E2.5, C57BL6/N embryos were either obtained from BlastoKit® (Charles River, Lyon, France) or collected from F1 female pregnant mice and cultured in KSOM media. 1 µM PD0325901 and 3 µM CHIR99021 (2i) were added to KSOM media in order to induce late blastocyst stage. FGF2 treatment was carried out at 100ng/ml in either KSOM from E2.5 or N2B27 from E4 stage. For analysis of Erk activity in late blastocysts, E2.5 embryos were cultured in KSOM/2i media for 2 days to E4.5. For analysis of Erk activity, embryos were then transferred to N2B27 alone or in presence of FGF2 (100ng/ml) for 30 min and immediately fixed in PBS containing 4% paraformaldehyde (PFA) at room temperature.

Immunofluorescence staining of blastocysts was done described previously (Nashun et al., 2015). In short, blastocysts were washed with 1% BSA/PBS three times. The cells were permeabilized by incubating in 1% BSA/PBS containing 0.5% Triton X-100 for 30 min at room temperature, and, after washing with 1% BSA/PBS three times, incubated with primary antibodies in 1% BSA/PBS containing 0.1% Triton X-100 overnight at 4ºC, washed three times, and incubated for 1 hour in the dark with Alexa Fluor 488- and/or 568-conjugated IgG secondary antibody (dilution 1:300, Molecular probes) in the same buffer. DNA was stained for 10 minutes with 3 µg/ml 4,6-diamidimo-2-phenylindole (DAPI) and the cells were then mounted with Vectashield solution (Vector Laboratories) on glass slides. Alternatively, embryos were subjected to antigen retrieval by dehydration/rehydration steps with increasing methanol concentrations in 0.1% TritonX-100-PBS. After extensive washing, embryos were incubated with acetone for 20min, blocked in 10% donkey serum and stained as described above.

Infection of embryos with lentiviral particles containing Erk-KTR-GFP was carried out by making a small hole into the zona pellucida of 8-cell stage (B6CBAF1 x B6CBAF1) embryos using the XYClone kaser (Hamilton Thorne). Lentivirus was expelled from the injection needle under the control of an Eppendorf Femtojet into the perivitelline space. Injected embryos were cultured in KSOM/2i media until E4.5. The embryos were then treated with either FGF2 or FGF4 (at 100ng/ml) in the presence or absence of Mek1/2 inhibitor PD032591 (5μM) for 30min. Embryos were fixed and stained for immunofluorescence.

Chimeras were generated by injection of B6N6 mES cells stably expressing H2B-mCherry and treated with either FGF2 or FGF4 (100ng/ml) in N2B27 for 36h into E4.5 blastocysts of albino B6-B6(Cg)-$Tyr^{c\text{-}Brd}$. Around 10-12 mES cells were injected into the blastocoel cavity of 30-31 blastrocysts per experimental condition using standard microinjection procedures. Embryos were transferred to pseudo-pregnant recipient mice. Embryos were dissected at E9.5, fixed in 4% PFA and assessed for chimeric contribution of injected H2B-mCherry cells. For FGF4 treated cells: 31 blastocysts were injected, 12 were dissected and 0 gave rise to chimeras. For FGF2 treated cells: 30 blastocysts were injected, 10 were dissected and 3 gave rise to chimeras.

##### Gastruloid generation

Generation of gastruloids was carried as previously described (Beccari, L. *et al.* 2018). Brifely, E14 mouse embryonic stem cells grown in 2i LIF media were harvested and resuspended in N2B27 media in presence of FGF2 (100ng/ml) or FGF4 (100ng/ml). 300 cells were plated in 40ul volume per well of low-adherence, round-bottom 96-well plate. Cells were incubated at 37 °C and 48h, cell aggregates were stimulated for 24h with 150ul of 3uM CHIR in N2B27. Thereafter media was replaced with N2B27 every 24h for seven days.

##### Clonogenicity assays

Following treatment, ES cells were seeded at 600 cells per well in six-well plates coated with 0.1% gelatine and cultured for 6 days in 2iLIF media. Cells were washed with PBS, fixed and stained for ALP (Sigma, 86R-1KT). Individual wells were imaged by SZX16 Wide Zoom Stereo Microscope and number of colonies were determined using ImageJ software.

#### Immunofluorescence and live cell imagining

Cells were washed in cold PBS and fixed with 4% paraformaldehyde in PBS for 10 min. Subsequently, cells were washed 3x in PBS, permeabilised with 0.3% Triton X-100 in PBS for 5 min and washed 3x in PBS. Samples were blocked with 10%FBS/3%BSA in PBS for 1h and incubated with the primary antibody at 4°C overnight. This was followed by 3x washes with 1% Tween in PBS and incubated with the secondary antibody at room temperature for 2 h. DAPI staining was used for nuclear staining and cell identification for image analysis purposes.

Imaging was performed on either ScanR, a fully motorized and automated inverted epifluorescence microscope system IX83 (Olympus), on an IncuCyte Zoom® (Essen BioScience) or on a Leica TCS SP8 confocal microscope (for embryo experiments) all equipped with temperature, humidity and $CO_2$ levels control.

ScanR images were typically acquired with a 20x plan (UCPLFLN) fluorescence objective (NA 0.7) and a sCMOS (Orca Flash 4.0, Hamamatsu) camera. LED-based illumination (SpectraX LED, Lumenco) was used for excitation. Excitation (ex) and emission (em) filters were as follows: DAPI ex: 391/20nm, em: 440/521/607/700nm; CFP ex: 438/24nm, em: 460-510nm, GFP/Alexa-488 ex: 474/27nm, em: 440/521/607/700nm; YFP ex: 509/22nm, em: 515-560nm, mCherry ex: 554/23nm, em: 440/521/607/700nm and Alexa-647 ex: 650/13nm, em: 690/50nm. IncuCyte Zoom images were acquired with a 20x plan fluorescence objectives and a CCD camera. Fluorescence excitation (ex) and emission (em) filters were as follows: Green channel ex: 440-480nm, em: 504-544nm; Red channel ex: 565-605nm em: 625-705nm.

#### Imaging of mouse gastruloids

Gastruloids were mounted in agarose 1% (SERVA 11384.01) by mean of glass capillary with inner diameter of 1.15 µm. Imaging was performed using a Bruker-Luxendo MuVi-SPIM light-sheet microscope equipped with two 20x objectives for detection (Olympus XLUMPLFLN 20XW NA 1.0), two 10x objectives for illumination (Nikon CFI Plan Fluor 10X W NA 0.3) and two CMOS cameras (Hamamatsu Orca Flash 4.0 v3). Image volumes were acquired using the following combination of laser lines and filters: 1) 405 nm and BP 418-462; 2) 488 nm and BP 499-569; 3) 594 nm and BP 610-651; z-step between planes has been set to 1 µm and a magnification changer was used giving a final magnification of 16.6x and a voxel size of 0.392 x 0.329 x 1 µm. Thickness of the light-sheet was 4.1 µm and the “line mode” (virtual confocal slit of 20 pixels) has been used to increase the image contrast. For samples bigger than the field of view tiling and stitching of multiple volume have been performed.

Final image volumes are the result of the fusion of the 2 opposite camera views. Registration and fusion of volumes was performed using the Luxendo processing software (GUI 2.9.0) and the FIJI plugins BigStitcher and BigDataProcessor. Data was downsampled two times for the video editing purpose

**Western blotting analysis**

Following stimulation, cells were washed in ice-cold PBS and lysed in ice-cold radioimmunoprecipitation assay buffer [50 mM tris(hydroxymethyl)aminomethane (Tris)-hydrochloric acid (HCl), pH 7.5, 150 mM sodium chloride (NaCl), 1% Nonidet P-40, 0.1% Sodium dodecyl sulphate (SDS), 0.5% sodium deoxycholate] containing phosphatase and protease inhibitor cocktails (Sigma). Cell lysates were run on SDS-PAGE and proteins were transferred onto PVDF membranes and incubated overnight at 4°C with primary and 1 hour at room temperature with secondary antibody. Proteins visualized and photographed using ECL Prime detection reagent (GE Healthcare) and ImageQuant LAS 4000 analysis System (GE Healthcare).

**In situ hybridization**

Mouse uterus samples were dissected, fixed in 10% neutral buffered formalin for 24 hours at room temperature, processed and embedded in paraffin. Sections were cut at 3 µm and mounted into SuperFrost PlusTM slides. RNAscope was performed according with the manufacturer’s instructions, using the RNAscope® 2.5 HD Assay – Brown Reagent kit (Bio-Techne, 322300). The tissue sections were pre-treated with RNAscope® Target Retrieval for 15 minutes and RNAscope Protease Plus for 27 minutes. The mRNA quality of the samples was assessed with the RNAscope Positive Control Probe – Mm-Ppib (Bio-Techne, 313911) and the RNAscope Negative Control Probe – DapB (Bio-

Techne, 310043), and Fgf2 signal was identified using RNAscope Probe – Mm-Fgf2 (Bio-Techne, 316851). The slides were counterstained with Harris Haematoxilin, dehydrated, cleared and mounted in a Sakura Tissue-Tek Prisma auto stainer.

**Proteomics and phospho-proteomics analysis**

**Sample processing**

Cellular lysates containing 550µg of protein in 8M urea, 100mM TRIS/HCL buffer (pH8.5) were loaded onto Microcon 30kD centrifugal filters (Merck Millipore, MRCF0R030). Samples were then digested using a Filter Aided Sample Preparation (FASP) protocol (Wiśniewski et al., 2009). Briefly, samples were concentrated on the filter unit through centrifugation (12,000g) and then buffer exchanged using sequential washing and centrifugation with 8M Urea buffer. Proteins were then reduced and alkylated sequentially with 10mM Dithiothreitol and 50mM Iodoacetamide (in 8m urea buffer) respectively. Samples were then further buffer exchanged to remove salts using sequential washing with 50mM ammonium bicarbonate (AmBic). Trypsin Gold (Promega, V5280) was added to the samples in 50mM AmBic to an approximate 1:50, protease:protein ratio. Digestions were incubated overnight (17h) at 37$^{o}$C. Digest extracts were recovered from FASP filters via centrifugation and acidified with 1% trifluoroacetic acid (TFA). Final digest extracts were then normalised to the same total volume with 1% TFA. For total protein analysis, volumes equivalent to 50µg of protein were desalted using Glygen C18 spin tips (Glygen Corp, TT2C18.96) and peptides eluted with 60% ACN + 0.1% formic acid (FA). Eluents were then dried using a centrifugal vacuum drier. The remaining sample volumes were de-salted using solid phase extraction with OASIS HLB 10mg cartridges (Waters, 186000383) according to the manufacturer's instructions.

Peptides were eluted with 1M glycolic acid in 80% acetonitrile (ACN), 5% TFA and phospho-enriched using a $TiO_2$ based method (Casado et al., 2014). Briefly, eluents were adjusted to 1mL with 1M glycolic acid solution and then incubated with 25mg of $TiO_2$ (50% slurry in 1% TFA) for 5 minutes at room temperature. After 5 minutes of incubation with mixing, the $TiO_2$/peptide mixture was packed into empty spin tips by centrifugation. The $TiO_2$ layer was then sequentially washed with 1M glycolic acid solution, 100mM ammonium acetate in 25% ACN and 10% ACN. Phosphopeptides were eluted with 4 sequential additions of a 5% $NH_4OH$ solution. Eluents were clarified with centrifugation (18,000g) for 2 minutes and clear supernatant transferred to new tubes. Samples were snap frozen on dry ice and then dried by vacuum centrifugation.

#### Liquid chromatography-tandem mass spectrometry (LC-MS/MS) analysis

Phospho-enriched samples and desalted total protein digests were redissolved in 0.1% TFA by shaking (1200rpm) for 30min and sonication on an ultrasonic water bath for 10min, followed by centrifugation (20,800g, 5°C) for 10min. LC-MS/MS analysis was carried out in technical duplicates (750ng on column for total protein samples) and separation was performed using an Ultimate 3000 RSLC nano liquid chromatography system (Thermo Scientific) coupled to a QE mass spectrometer (Thermo Scientific) via an EASY spray source (Thermo Scientific). For LC-MS/MS analysis total protein and phosphopeptide solutions were injected and loaded onto a trap column (Acclaim PepMap 100 C18, 100μm × 2cm) for desalting and concentration at 8μL/min in 2% acetonitrile, 0.1% TFA. Peptides were then eluted on-line to an analytical column (Acclaim Pepmap RSLC C18, 75μm × 50cm) at a flow rate of 250nL/min. Peptides were separated using a 120 minute gradient, 4-25% of buffer B for 90 minutes followed by 25-45% buffer B for another 30 minutes (composition of buffer B – 80% acetonitrile, 0.1% FA) and subsequent column conditioning and equilibration. Eluted peptides were analysed by the mass spectrometer operating in positive polarity using a data-dependent acquisition mode. Ions for fragmentation were determined from an initial MS1 survey scan at 70,000 resolution, followed by HCD (Higher Energy Collision Induced Dissociation) of the top 12 most abundant ions at 17,500 resolution. MS1 and MS2 scan AGC targets were set to 3e6 and 5e4 for maximum injection times of 50ms and 50ms respectively. A survey scan m/z range of 400 – 1800 was used, normalised collision energy set to 27%, charge exclusion enabled with unassigned and +1 charge states rejected and a minimal AGC target of 1e3.

#### Kinase activity predictions and proteomics data analysis

Missing mass-spectrometry detection values in the phosphoproteomics data-set were flagged and treated as missing information in any of the downstream calculations. The data-set was then reduced to consider only single phosphorylated phosphopeptides that mapped unambiguously to a single protein. Phosphopeptides mapping to the same protein Serine / Threonine / Tyrosine residue were averaged. The data-set was log2 transformed and z-score centred. Technical replicates were averaged and biological replicates were utilised to assess reproducibility by performing pearson correlation analysis between samples within same experimental conditions. Differential phosphorylation analysis was performed with Limma R package (v3.32.10) comparing conditions: N2B27 + FGF2 treated, Control (N2B27 only), and FGF2_Control for both time points, 15 and 60 mins. Statistical assessment of the mean phosphorylation log fold-changes was estimated using moderated t-tests using the different biological replicates.

Kinase/substrate (K/S) interaction network for *Mus musculus* was assembled from PhosphositePlus. This network was assembled considering *Mus musculus* specific K/S interactions and also K/S interactions identified in human, which were previously mapped using ensembl orthologue mapping. This comprised a total of 7,927 interactions, 322 unique kinases and 5,789 unique phosphorylation-sites. Kinase activity scores were *in silico* estimated using a two-sided z-test, similarly to PMID: 28200105, for kinases that contained at least 1 target phosphorylation-site, from the K/P interaction network, measured.

#### qRT-PCR analysis

Total RNA was extracted using TRI reagent (Life Technologies) and MaXtract High Density tubes (Qiagen) according to the manufacturer's instructions. To test mRNA expression in different mouse tissues, uterus, liver and thymus were collected from sacrificed E4.5 pregnant mice and from non-pregnant mice. Sample tissues were finely cut with a scalpel, mix in TRI reagent and intermittently vortexed for 30min. Tissue samples were subsequently frozen at -80C before RNA extraction. RNA purity and quantity was assessed by nanodrop (Fisher Scientific) (A260/A280 1.8-2 was considered suitable for further analysis), possible contaminating DNA was removed and cDNA prepared from 1 μg RNA using QuantiTect Reverse Transcription Kit (Qiagen) according to the manufacturer's instructions. qRT-PCR was performed on a Bio-Rad CFX96 real-time detection system using iTaq SYBR Green qPCR master mix (Bio-Rad) and primer pairs as listed in table 1. PCR conditions consisted of 1 cycle of 95 °C for 3 min and 40 cycles of 95 °C for 5 sec and 60 °C for 30 sec. Both GAPDH and glucuronidase beta (GUSβ) were used as housekeeping genes.

**Table 2.** Primers for qPCR used in this study

| Gene name | Sequence (5' to 3') | Sequence (5' to 3') |
|---|---|---|
| Cdx2 | CAGGAGGAAAAGTGAGCTGG | TCTTGATTTTCCTCTCCTTGGC |
| Dax1 | CGTGCTCTTTAACCCAGACC | CCGGATGTGCTCAGTAAGG |
| Dnmt1 | GAGCGTTGTGGTGGATGAC | GGGAGGTCAGACATGGTGT |
| Dnmt3a | CCTGCAATGACCTCTCCATT | CAGGAGGCGGTAGAACTCAA |
| Dnmt3b | TGGTGATTGGTGGAAGCC | AATGGACGGTTGTCGCC |
| Dnmt3l | CGCATAGCATTCTGGTAGTCTCTG | ATGGACAATCTGCTGCTGACTG |
| Elf3 | TGGGGCCAGAAGAAGAAGAA | CCATCCACCCGTTCCAGG |
| Esrrb | GGCGTTCTTCAAGAGAACCA | CCCACTTTGAGGCATTTCAT |
| FGF2 | CAAGGGAGTGTGTGCCAAC | CCAGTCGTTCAAAGAAGAAACA |
| Foxa2 | TGAAGATGGAAGGGCACGAG | CATTCATCCCCAGGCCGG |
| Gapdh | CCGGTGCTGAGTATGTCGT | GGGCGGAGATGATGACCC |
| Gata6 | AGCAAGATGAATGGCCTCAG | GGTTGTGGTGTGACAGTTGG |
| Gbx2 | GGCACCTCCTAGATGTGGAC | AAAACACTGCAGCTGAGATCC |
| Hoxb1 | CGGACCTTCGACTGGATGAA | GTGAAGTTTGTGCGGAGACC |
| Klf2 | CTAAAGGCGCATCTGCGTA | TAGTGGCGGGTAAGCTCGT |
| Klf4 | CGGGAAGGGAGAAGACACT | GAGTTCCTCACGCCAACG |
| Klf5 | CCGGAGACGATCTGAAACAC | CAGATACTTCTCCATTTCACATCTTG |
| Mb3 | AGAAGAACCCTGGTGTGTGG | TGTACCAGCTCCTCCTGCTT |
| Mi2b | GCCAATGCAGTCCTACACAA | TGTAACCTCACAGCGACTGG |
| Nanog | TACCTCAGCCTCCAGCAGAT | GCTTGCACTTCATCCTTTGG |
| Pax3 | CTGTCACAGGCTACCAGTATG | GAAACAGGGCTCCAAGTGG |
| Pou3f3 | CCATGCCTAAAGATTCCGCC | TTGGTGCTGTCTTTCCACAC |
| Pou5f1 (Oct4) | GAGGAAGCCGACAACAATGA | CCTCACACGGTTCTCAATGC |
| Prdm14 | CAGCCTGAACAAGCACATGA | TGTGTTCGGAGTATGCTGGA |
| Rex1 | GCGCTATCTCAACCTGTTCA | GGTCATACGCCACCTTTTCC |
| Sall4 | GAAGCCCCAGCACATCAAC | CTGAGGCTTCATCGCAGTT |
| Sox1 | GGCTTCGGAGGACAAAAGAC | AAGAGCTGGCGGGAAGTAAA |
| Sox17 | GTGTGGGCCAAAGACGAAC | TCAACGCCTTCCAAGACTTG |
| Sox2 | GACCGTTTTCGTGGTCTTGT | CGATATCAACCTGCATGGAC |
| Stat3 | GTCCTTTTCCACCCAAGTGA | TATCTTGGCCCTTTGGAATG |
| Tbx3 | TTGCAAAGGGTTTTCGAGAC | TGCAGTGTGAGCTGCTTTCT |
| Tcf3 | CTGAGCAGCCCGTACCTCT | AGGGGCCATTTCATCTGTAG |
| Tet1 | TTCACAACATGCACAACGGA | CAGGACGTGGAGTTGTTCATC |
| Tet2 | GCTCATTCCCACAGAGACCA | CTCAGGCTTAGCTCCGACTT |
| Tet3 | CACGCCAGAGAAGATCAAGC | CTTCAGGCTTTGCTGGGAC |
| Tfcp2l1 | GGGGACTACTCGGAGCATCT | TTCCGATCAGCTCCCTTG |

#### Bisulfite Sequencing

DNA was isolated using the Qiagen Blood and Cell DNA Mini kit and bisulfite conversion was carried out using the Qiagen EpiTect Kit according to manufacturer's instructions. Converted DNA was PCR amplified using KAPA HiFi HotStart Uracil+ ReadyMix kit and previously described primer sets (Table 2). PCR conditions were 94°C for 5 min, followed by 30 cycles of 98°C 20 sec, 57°C 30 sec, 72°C 30 sec and final extension of 1 min at 72°C. PCR products were gel purified using Qiagen gel extraction kit and cloned into a pGEM-T Easy vector (Promega) after A-tailing procedure as described by manufacturer. Positive colonies were subjected to colony PCR and amplified products were purified and sequenced. The methylation status of target sequence was determined by quantification tool for methylation analysis (QUMA) software.

**Table 3**: Primers used in this study for bisulfite sequencing

| Region | Sequence | Size |
|---|---|---|
| Kcnq1ot1 Fw in | GGT TAG AAG TAG AGG TGA TT | 206 |
| Kcnq1ot1 Fw out | GTG TGA TTT TAT TTG GAG AG | |
| Kcnq1ot1 Rev in | CAA AAC CAC CCC TAC TTC TAT | |
| Kcnq1ot1 Rev out | CCA CTC ACT ACC TTA ATA CTA ACC AC | |
| LINE1 Fw | TAGGAAATTAGTTTGAATAGGTGAGAGGT | 282 |
| LINE1 Rev | TCAAACACTATATTACTTTAACAATTCCCA | |
| Tcl1-R3 Fw in | GTTAGTAGGTGGGGTAGTAGGGATATG | 297 |
| Tcl1-R3 Fw out | GAAATAGGAGGGTTAGGGAGATTTTAGATG | |
| Tcl1-R3 Rev in | CTTCACACACAATCTCAACAATTTACTTTTAA | |
| Tcl1-R3 Rev out | TTTCTTTTAAACACCAACATTAAAACCCATAC | |

#### Quantification and statistical analysis

##### Statistical analysis

Statistical comparisons between means were made by one-way ANOVA, Mann-Whitney and Kolmogorov–Smirnov (KS)-tests (non-parametric tests) and using student T-test to evaluate significance between different experimental conditions. A *p*-value of less than 0.05 was considered statistically significant.

#### Image Analysis

Image analysis was carried out using ScanR software, Imaris Bitplane 9.2.1, ImageJ or custom built Matlab scripts. Fluorescent images of the mES nucleus were segmented to generate nuclear masks. Segmented nuclear masks were then used to quantify the intensity of reporter proteins or biosensors from the background subtracted fluorescent channel images. To quantify the nuclear cytoplasmic ratio for each nucleus, mean fluorescent intensity of the 3-pixel width ring around nucleus was divided by the mean fluorescent intensity of pixels within the nucleus. In time course experiments the population average was calculated as mean of the nuclear cytoplasmic ratio in all cells and was plotted over time to visualize the dynamics. Embryo images were segmented manually by drawing ROI in ImageJ and intensities quantified directly after background subtraction.

#### Proteomics raw data processing

Data was processed using the MaxQuant software platform (v1.5.8.3), with database searches carried out by the in-built Andromeda search engine against the Swissprot *mus musculus* database (version 20170202, number of entries: 16,844). A reverse decoy database approach was used at a 1% false discovery rate (FDR) for peptide spectrum matches. Search parameters included: maximum missed cleavages set to 2, fixed modification of cysteine carbamidomethylation and variable modifications of methionine oxidation, protein N-terminal acetylation and serine, threonine, tyrosine phosphorylation for the phospho-enriched data-set. For the total protein samples fixed modification of cysteine carbamidomethylation and variable modifications of methionine oxidation, protein N-terminal acetylation, asparagine deamidation and cyclization of N-terminal glutamine to pyroglutamate. Label-free quantification was enabled with an LFQ minimum ratio count of 2. 'Match between runs' function was used with match and alignment time limits of 1 and 20 minutes respectively.

**RNA-Seq Analysis and batch effect correction**

Identification of genes changing across time was performed using the DESeq2 version 1.2.10 Bioconductor library (Love et al., 2014). Differential genes were selected by fitting an LRT model across the time-points. Genes differential at any one time point were selected using a 0.05 false-discovery rate (FDR) threshold and an absolute log2 fold change > 2 at any one time-point when compared to time zero.

RNAseq from mouse embryo data from E3.5, E4.5 and E5.5 and mES cells treated with 2iLIF was acquired from Broviak et al. 2015 RNA-seq data, obtained from the ENA (https://www.ebi.ac.uk/ena/data/view/PRJEB7393). Similarly to the RNA-seq data acquired in this study, data was processed with the nextflow BABS-RNAseq pipeline which includes adapter trimming with Cutadapt (DOI: 10.14806/ej.17.1.200, https://cutadapt.readthedocs.io/en/stable/) and read alignment and quantification with STAR (DOI: 10.1093/bioinformatics/bts635 , https://github.com/alexdobin/STAR) and RSEM (DOI: 10.1186/1471-2105-12-323 , https://github.com/deweylab/RSEM). We aligned the data against the mouse genome (version: GRCm38 ensembl release 89).

After combining the quantification results from both experiments, we removed genes that were detected with less than 1 read across all samples and applied variance stabilizing transformations with the vst() function from the DESeq2 R package (version 1.24.0, DOI: 10.1186/s13059-014-0550-8). To enable the comparison between the Boroviak et al. 2015 and the current dataset, we removed batch effects using the removeBatchEffect() function from the limma package (version 3.40.2, DOI: 10.1093/nar/gkv007) using R.

The 500 differentially expressed genes upon FGF2 treatment over time with the lowest adjusted p-value were used for visualization of the first two principal components of the combined Gharibi et al. and Boroviak et al. data sets using the plotPCA() function from the DESeq2 R package (version 1.24.0, DOI: 10.1186/s13059-014-0550-8) and ggplot2 (version 3.2.1, Wickham H (2016). ggplot2: Elegant Graphics for Data Analysis. Springer-Verlag New York. ISBN 978-3-319-24277-4, https://ggplot2.tidyverse.org.).

**Data and code availability**

All proteomics raw data that supports the findings of this study have been deposited in PRIDE repository hosted by the EBI-EMBL with the dataset identifier with identifier number PXD015851.

Raw sequencing data sets (RNA-seq) that support the findings of this study have been deposited to GEO repository (hosted by NCBI) under the superseries identifier GSE156672.

**Supplemental Movies**

**Movie S1. FGF2 induces a transient activation of Erk1/2 in mES cells.** Erk activity in real time of mES cells expressing Erk-KTR sensor and treated with N2B27 in presence (right) or absence (left) of FGF2. Cells were imaged every 15min for duration of 24h.

**Movie S2. FGF2 and FGF4 induce different proportions of embryonic lineages.** (Left) Light-sheet 3D reconstruction showing mouse gastruloids at 168h after FGF2 stimulation for 48h stained with primitive streak/mesoderm marker Brachyury (green), endoderm marker (FOXA2) and DAPI (grey). (Right) Light-sheet 3D reconstruction showing mouse gastruloids at 168h after FGF4 stimulation for 48h stained with primitive streak/mesoderm marker Brachyury (green), endoderm marker (FOXA2) and DAPI (grey).
